# One Scaffold, Many Targets: Fully Modified RNA Nanostructures for Multi-Gene Silencing Beyond the Liver

**DOI:** 10.64898/2026.10.08.757482

**Authors:** Hassan H. Fakih, Ashley Summers, Qi Tang, Rosemary Gagnon, Brianna Bramato, Raymond Furgal, Katherine Y. Gross, Jana H. Fakih, Dimas Echeverria, Hanadi Sleiman, Anastasia Khvorova

## Abstract

Therapeutic siRNAs are a major clinical modality for hepatic gene silencing, but efficient extrahepatic delivery and multi-gene modulation remain significant barriers. Nucleic acid nanotechnology offers programmable assembly, but conventional structures suffer from enzymatic instability *in vivo*. Here we report fully chemically modified RNA nanostructures (FMRNs)—a vehicle-free platform entirely composed of stabilized therapeutic nucleic acids that integrate structural precision with metabolic durability. Following systemic administration, FMRNs exhibit enhanced pharmacokinetics and extrahepatic accumulation, with *in vivo* behavior influenced greatly by hydrodynamic size rather than molecular weight. Using triangular and square geometries, we demonstrate programmable control over siRNA stoichiometry to achieve sustained, simultaneous silencing of up to four genes from a single entity. Nanostructure assembly alone drives productive delivery, especially to heart and muscle, matching lipophilic conjugation. Furthermore, cerebrospinal fluid administration reveals that larger FMRNs improve central nervous system distribution and retention revealing physicochemical parameters that impact brain exposure. FMRNs establish a modular and effective platform combining improved pharmacokinetics, extrahepatic delivery, and multi-gene silencing for complex diseases.

## INTRODUCTION

Oligonucleotide therapeutics, including small interfering RNA (siRNA), have emerged as a new class of medicines due to their ability to precisely modulate disease-causing genes with multi-month efficacy. ^1,2^ To date, over twenty oligonucleotide therapeutics, including multiple siRNA-based drugs, have received regulatory approval. ^3^ This clinical success follows decades of advancements in chemical modification strategies that confer robust metabolic stability, safety and sustained gene silencing. ^1,3^ Despite these advances, currently approved siRNA drugs remain restricted to the liver, a consequence of the unparalleled success of the trivalent N-acetylgalactosamine (GalNAc) conjugate, which mediates efficient delivery to hepatocytes.^1,3^ To unlock the full therapeutic scope of siRNAs, focus has shifted toward technologies that can extend gene silencing to extrahepatic tissues, including the central nervous system (CNS), heart, and skeletal muscle. ^3,4^

Current extrahepatic delivery efforts primarily leverage two distinct strategies: molecular-scale conjugates and nanotechnology systems.^3,4^ Conjugates (e.g., lipophilic moieties, peptides, or antibodies) offer structural homogeneity, defined stoichiometry as single entities, and predictable pharmacokinetics.^1,3–5^ However, they are often limited by low delivery efficiency per conjugate, restricted payload capacity, and tedious medicinal chemistry for some.^3,6,7^ Notably, divalent siRNAs (di-siRNAs) — in which two siRNA duplexes are directly linked to one another, rather than to an exogenous chemical conjugate — demonstrate that an entirely RNA-based, unimolecular architecture can itself confer conjugate-like pharmacokinetic behavior, without requiring a separate delivery moiety. This approach has proven particularly effective in the CNS, where di-siRNAs achieve enhanced potency and durability^8^, suggesting that engineering size and structure directly into the RNA itself, rather than through external conjugation, may offer a general, chemically self-contained strategy for tuning pharmacokinetics and pharmacodynamics. Conversely, nanotechnology platforms, like lipid nanoparticles (LNPs), encapsulate dense RNA payloads but are burdened by complex manufacturing, batch-to-batch variability, and potential vehicle toxicity.^9,10^ Consequently, there remains a significant need for a delivery platform that combines the molecular precision of conjugates with the high-capacity payload of nanoparticles.

As siRNA therapeutics mature toward tackling complex polygenic diseases (e.g., cancer^11^, neurodegenerative^12^ and inflammatory diseases^13,14^), multi-target or multi-pathway modulation is emerging as an essential strategy.^11,14^ Clinically advanced multi-targeting approaches, such as Comanche Biopharma’s fixed-dose combination of two lipid-conjugated siRNAs targeting distinct *sFLT1* isoforms, rely on physical mixtures of separate compounds that require matched pharmacokinetics.^15,16^. Furthermore, in complex disorders such as neurological diseases (e.g., Huntington’s and Alzhimer’s disease), cardiometabolic disorders, cancer, and autoimmune diseases, simultaneous multi-target modulation is key to maximize efficacy against disease and address malignancy along with resulting dysfunctions.^17–19^ Additionally, the recent reportings on genetic compensation highlight the need for multi-targeting to ensure kockdown and biological effects. ^20–22^ Structural nucleic acid nanoparticles (NANPs) offer a promising frontier to address this need, leveraging base-pairing rules to self-assemble “vehicle-free” architectures that afford inherent biocompatibility and absolute stoichiometric control from a single RNA-based material.^23,24^ By eliminating the need for complex mixtures, NANPs provide a unique, single-material framework capable of delivering multiple distinct oligonucleotide therapeutics with precise spatial control.^23–27^ Beyond enabling this stoichiometric and spatial control, an all-RNA architecture of this kind may also confer the improved pharmacokinetic and pharmacodynamic behavior already established for divalent siRNAs, extending that same unimolecular, conjugate-free design principle to higher-order, multi-target structures which become synthetically challenging to achieve through direct linear linkage alone.^8,28^ A diverse array of two- and three-dimensional NANPs designs have been reported, demonstrating the modularity of this approach and providing a unique platform to systematically interrogate how size, shape, and functionalization dictate *in vivo* behavior.^23,26,29–32^ Prior work has used NANPs to deliver siRNAs and antisense oligonucleotides (ASOs), with examples in cancer and immunotherapy settings.^33,34^

However, the *in vivo* utility of NANPs has been hindered by limited efficacy and durability, likely due to lack of robust biological stability.^24^ While innovation in structural design has accelerated, the field has largely relied on unmodified or minimally stabilized scaffolds, or has restricted chemical modifications solely to the therapeutic payload.^9,26,29,35–38^ Remarkably, most reported structures investigated for drug delivery remain constrained to phosphorothioate (PS) backbone alterations, with many more recently utilized modifications unused.^24^ This highlights a significant gap between the nanotechnology and medicinal oligonucleotide chemistry fields, which possesses a clinically proven toolkit of modifications —such as patterned 2ʹ-fluoro (2ʹ-F) and 2ʹ-O-methyl (2ʹ-OMe) sugar modifications and 5ʹ-vinylphosphonate (5ʹ-VP)— that have been proven to improve stability and potency by orders of magnitude,^1,39^ leaving their true multi-gene silencing potential and comparative performance against state-of-the-art technologies (e.g., lipid conjugates and multivalent architectures) completely unexplored.^9,10,24^

Here, we bridge the nanotechnology and oligonucleotide therapeutics fields by engineering fully chemically modified RNA nanostructures (FMRNs): “simple yet stable” architectures constructed *entirely* from clinical-grade chemically-stabilized nucleotides. FMRNs capitalize on the structural advantages of nanoparticles—enhanced pharmacokinetics and multi-target capacity—while attaining the stability and potency of chemically mature oligonucleotide therapeutics. We systematically describe the design, assembly, and characterization of triangular and square FMRNs carrying three and four distinct siRNAs, respectively. FMRNs maintain structural integrity in biological media and exhibit pharmacokinetics comparable to a lipid-conjugated siRNA. Crucially, systemic administration of FMRNs enables potent, simultaneous silencing of four distinct genes in extrahepatic tissues (including heart and muscle) sustained for at least one month. Following CNS delivery, FMRNs achieve safe, effective silencing of four targets for up to six months, with geometries exhibiting distinct local tissue distribution trends across brain regions. This establishes FMRNs as a robust, modular platform for multi-oligotherapeutic delivery for the potential treatment of complex polygenic diseases.

## RESULTS

### Design and optimization of FMRN assembly conditions

To evaluate the feasibility of constructing FMRNs, we designed two polygonal architectures: a triangle (T-FMRN) and a square (S-FMRN) that assemble from fully-modified “Arm” strands (Fig. 1a). These geometries were selected for their structural simplicity, requiring only three or four Arm strands, respectively, and featuring shared Arm strand motifs to facilitate comparative analysis of geometry-dependent biological activity (Extended Fig. 1). The Arm strands used for the initial design of FMRN assembly are 58-nucleotides (nt) long and consist of three functional domains: (i) two 15-nt scaffolding regions for inter-Arm strand hybridization; (ii) a 20-nt therapeutic-binding domain designed to accommodate ASOs or siRNAs; and (iii) a 4-nt flexible vertex to facilitate precise folding that remains single-stranded (Extended Fig. 1a). Every nucleotide in each Arm strand was chemically modified with either 2ʹ-OMe or 2ʹ-F sugar modifications, and PS backbone linkages were added at select positions, mirroring the modification patterns utilized in clinically approved siRNA drugs. Full details of the sequences are provided in Supplementary Table 1.

**Figure 1:**
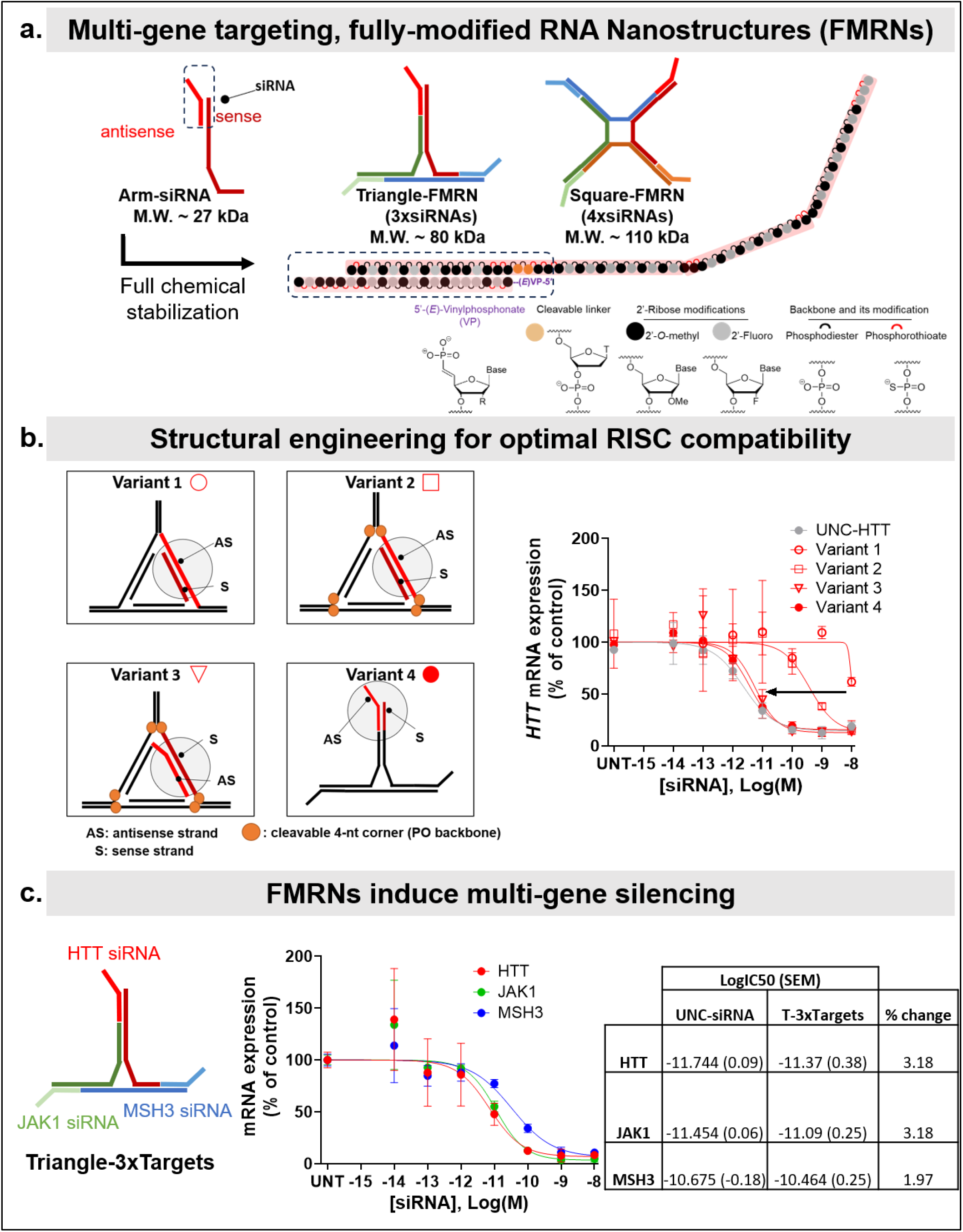
Design and systematic optimization of FMRNs for siRNA placement enables maximum efficacy, and mutli-gene targeting. **a.** Cartoon schematic of polygon-based fully modified RNA nanostructures (FMRNs). They are designed from various “Arms” that assemble into a triangular “Triangle” or quadrangular “Square” nanostructures. Nucleotide-based resolution schematic shows that each nucleotide in all strands are fully chemically modified with standard chemistry used in therapeutic siRNAs : 2’Fluoro (2’F), 2’ O-methyl (2’OMe), or Phosphorothioate (PS), and 5’ Vinyl Phosphonate at the 5’ terminus of the antisense strand. **b.** T-FMRN design variation for the placement of siRNA component, to optimize for RISC compatibility and maximum siRNA efficacy. Version 1 positions the antisense (AS) strand as part of the polygon arm. Version 2 introduces cleavable corners (phosphodiester backbone instead of phosphorothioate) that facilitate cleavage and release of the siRNA. Version 3 swaps position of AS to be hybridized with keeping the cleavable corners. Version 4 positions the siRNA at the extremity of the FMRN. Dose-dependent siRNA efficacy *in vitro* targeting *Htt* mRNA reveals improved potency from version 1 to 4, to levels comparable to that of control unconjugated (UNC)-HTT siRNA. **c.** Multi-gene targeting capabilities of T-FMRN confirmed *in vitro,* with nanostructure carrying siRNAs targeting HTT, JAK1 and MSH3 mRNAs simultanously. Potency is comparable to control UNC-siRNA, targeting these genes, with comparable IC50s. **c.** Multi-modality functionalization of T-FMRN, by delivering one ASO targeting *HTT* and two siRNAs targeting *JAK1* and *MECP2* mRNAs. Dose-dependent siRNA efficacy *in vitro* against each target is observed. (all *in vitro* studies are conducted in HeLa cells, with siRNAs transfected at the indicated concentrations for 72h ; n=3/group; mRNA quantification via Quantigene 2.0 assay).

Initial assembly was performed in magnesium-containing tris buffer (7.6 mM Magnesium Chloride) using a conventional slow-cooling cycle (2h).^40,41^ Polyacrylamide gel electrophoresis (PAGE) revealed sharp, single bands for both T- and S-FMRNs, indicative of high-yield, monodisperse assembly (Extended Fig. 1b). To enable eventual *in vivo* utility, we optimized assembly in standard phosphate-buffered saline (PBS) typically used for oligonucleotide formulations and confirmed the quantitative loading of three or four siRNA duplexes into T- and S-FMRNs, respectively. We observed successful assembly and payload loading, as evidenced by distinct, single bands visualized by nucleic acid staining (Extended Fig. 1c).

To streamline structure assembly, we investigated whether the 2-hour annealing step could be eliminated, as full chemical modification significantly enhances thermodynamic stability^1^, and thus likely to improve hybridization kinetics-and FMRN assembly. Additionally, to assess suitability for high-dose *in vivo* administration, we evaluated assemblies at concentrations relevant for systemic and local delivery (100 µM siRNA equivalent, representing 33.34 µM T-3xsiRNA and 25 µM S-4xsiRNA carrying 3 or 4 copies of the same siRNA duplex, respectively). We observed that the rigorous 2-hour annealing process could be replaced with a rapid, 5-minute 95°C “quick-anneal”, the standard protocol for siRNA duplex preparation, without compromising structural integrity (Extended Fig. 1d). Both T- and S-FMRNs yielded well-defined, monodisperse bands with or without siRNA loading. These results suggest that the increased thermodynamic stability of fully modified oligonucleotides facilitates nanostructure assembly and potentially enables the development of structural architectures that would be challenging to achieve using conventional DNA-based nanostructures.

### Nanostructure architecture profoundly impacts siRNA efficacy

Following optimization of the assembly conditions, we sought to optimize the FMRN architectural parameters governing siRNA potency. We hypothesized that the full modification and structural rigidity of FMRNs (Fig. 1a) might impose steric constraints on the loading of the siRNA antisense strand into its effector complex (the RNA-induced silencing complex, or RISC) and on RISC activity. To systematically interrogate this structure-activity relationship, we evaluated four design iterations of the T-FMRN, varying the spatial orientation and connectivity of the siRNA payload (Fig. 1b, left). In Variant 1, the siRNA antisense strand was directly integrated into the Arm strand to maximize nuclease protection, albeit at the cost of high steric hindrance. Variant 2 involved the selective removal of PS modifications in the single-stranded vertices to facilitate siRNA release. Variant 3 used a “flipped” orientation, where the antisense strand was hybridized to the Arm strand rather than integrated into the Arm backbone. Variant 4 positioned the hybridized siRNA at the distal extremities of the FMRN arms to maximize accessibility, with keeping the vertices’ PS modifications. For each variant, siRNAs were designed using a previously validated sequence that silences the ubiquitously expressed huntingtin (*HTT*) gene.^8,42^

The *in vitro* potency of Variants 1-4 and a positive siRNA control (unconjugated, HTT-targeting siRNA, UNC-HTT) was assessed in *HeLa* cells, using a 7-point dose-response assay (Fig. 1b, right). While Variant 1 exhibited significantly attenuated silencing activity compared to control siRNA, Variant 2 recovered some of the efficacy yet significantly less compared to the siRNA control. Increasing accessibility of the antisense strand using a flipped orientation (Variant 3) or distal extremity placement (Variant 4) resulted in potencies similar to the positive siRNA control. Variant 4 was prioritized for subsequent investigations due to its optimal balance of potency and structural accessibility (Fig. 1b, left). Step-wise assembly confirming monodisperse for this variant construct for both T-FMRN and S-FMRN was also characterized via gel electrophoresis and provided in the supplementary data (Supplementary Fig. 1).

### Nanostructures enable multi-gene silencing

Leveraging the inherent valency of the FMRN architecture, we next investigated its capacity to simultaneously modulate multiple genes. We loaded T-FMRN (the optimal variant 4) with three distinct, previously validated siRNAs targeting *HTT* mRNA, janus kinase 1 (*JAK1)* mRNA, or mutS omolog 3 *(MSH3)* mRNA, and determined *in vitro* potencies using a 7-point dose-response assay (Fig. 1c, left). Upon transfection, we observed potent silencing of all three targets within the same cellular population. When comparing the IC50 for each T-FMRN siRNA to the respective unconjugated siRNA control (UNC-HTT, UNC-JAK1, and UNC-MSH3 delivered individually), no significant difference was observed. These results confirm the platform’s capacity for polygenic intervention without compromising individual siRNA efficacy (Fig. 1c, right).

To further evaluate the modularity of the FMRN delivery system, we loaded a heterogeneous therapeutic payload consisting of one ASO (targeting *HTT* mRNA) and two distinct siRNAs (targeting *JAK1* and Methyl-CpG binding protein 2, *MECP2* mRNAs) into T-FMRN, and measured *in vitro* potencies. This multi-modal architecture maintained potent activity across all three targets, highlighting the versatility of FMRNs in delivering diverse oligonucleotide classes for the treatment of multifaceted diseases (Supplementary Fig. 2).

### Full chemical modification is essential for metabolic stability, tissue accumulation, and silencing *in vivo*

Following identification of the optimal structural design (Variant 4), we set out to evaluate the stability of the fully chemically modified nanostructures. We compared the metabolic integrity of T-FMRN loaded with three *HTT*-targeting siRNAs (T-3xHTT-FMRN) to that of a control triangular architecture assembled from DNA stabilized with PS (T-3xHTT-DNA) (Fig. 2a), the most common modification for nanostructures.^23,24^ To isolate the effect of scaffold chemistry, T-DNA was designed with PS modifications at the 5ʹ and 3ʹ extremities of each arm and at the corners—providing a basal level of stabilization—while maintaining the siRNA payload in its fully modified form (detailed PS modifications provided in supplementary table 1). Characterization of the assembly of both constructs was confirmed via gel electrophoresis and dynamic light scattering (Supplementary Fig. 3a,b), demonstrating near-perfect assembly and similar sizes (T-DNA: ∼13 nm; T-FMRN: ∼12 nm). Following incubation of each architecture in cell culture media supplemented with 10% fetal bovine serum (FBS) rich in nucleases that degrade oligonucleotides, a time-course analysis was carried out via native gel electrophoresis (Fig. 2b, with gels provided in Supplementary Fig. 3c). The FMRN exhibited minimal degradation or disassembly for up to seven days, whereas the T-DNA structures underwent complete degradation within one hour, conferring the insufficiency of conventional PS-backbone stabilization strategy to confer metabolic stability (Fig. 2b).

**Figure 2:**
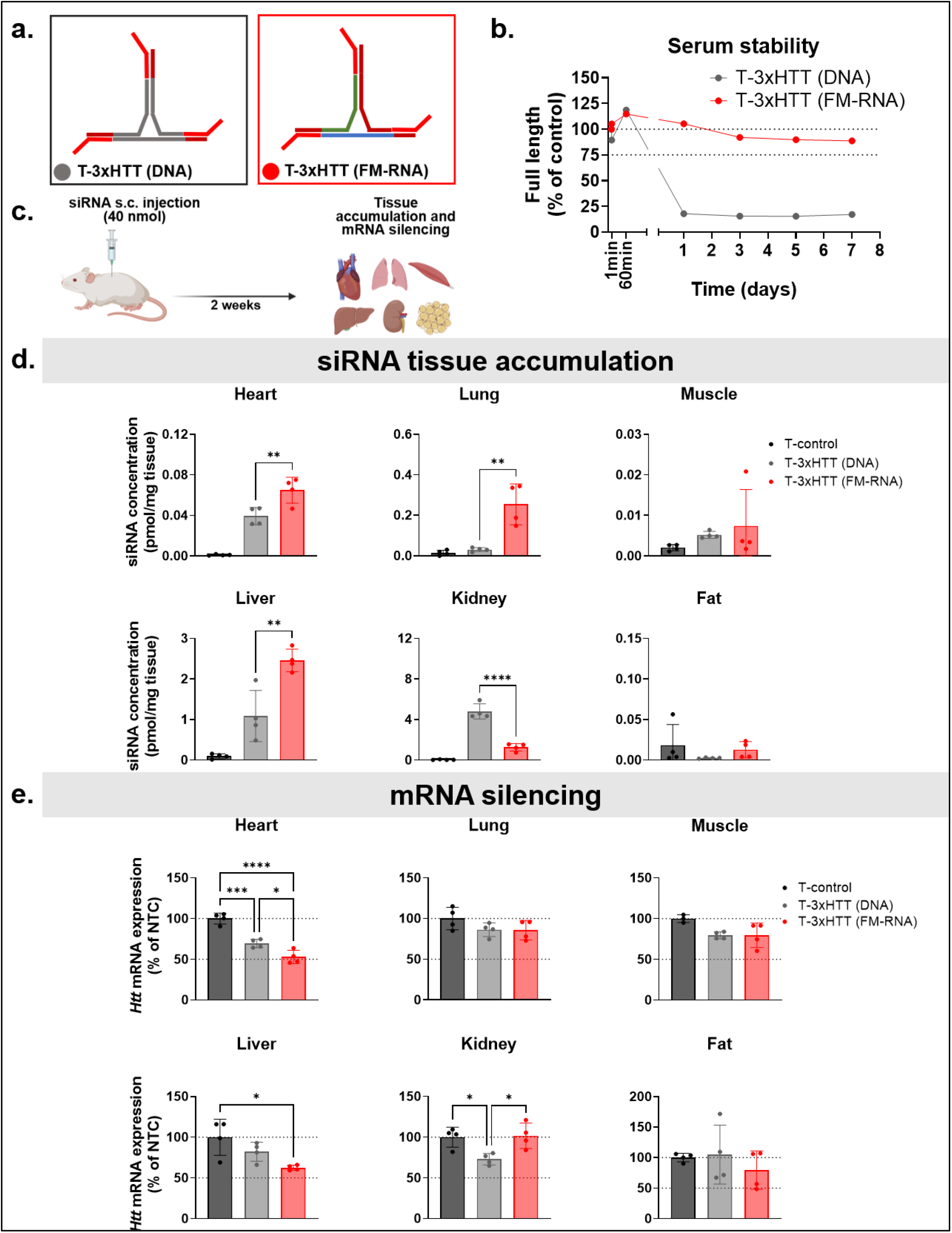
Fully chemically stabilized T-FMRN outperforms partially modified T-DNA in siRNA tissue accumulation and target silencing following systemic injection *in vivo*. **a.** Schematic of partially-modified DNA-based Triangle carrying modified siRNAs targeting HTT mRNA (T-3xHTT (DNA)) and Fully modified T-FMRN (T-3xHTT (FMRN)). **b.** Serum stability assay quantifying the stability of each of the structures following incubation in serum-containing cell culture media (10% FBS supplemented DMEM media) via quantification of intact band on native PAGE gel (provided in supplementary Figure 3). **c.** Scheme of *in vivo* study comparing both structures, with mice injected subcutaneously (s.c.) with T-3xHTT, T-3xHTT (DNA), or T-control (T-FMRN without any antisense strands) at a 40 nmol siRNA dose (equivalent to 20 mg/kg siRNA dose), with tissues assessed for siRNA tissue accumulation and mRNA silencing two-weeks post injection. **d.** siRNA Tissue accumulation and **e.** mRNA silencing two-weeks post s.c. injection, with siRNA activity compared to T-FMRN not loaded with antisense strands acting as non-targeting control. All study groups are *n* = 3-4 mice/group. Statistical analysis was performed in GraphPad Prism: one-way ANOVA analysis followed by Tukey’s multiple comparisons across all groups, with comparisons between T-3xHTT and T-3xHTT (DNA) only being shown for accumulation. ; non-significant comparisons are not shown, \**p* < .05, \*\**p* < .01, \*\*\**p* < .001, \*\*\*\**p* < .0001 (tissue accumulation measured by PNA assay; mRNA measured by the Quantigene 2.0 Assay; data represented as mean ± S.D.; dotted line on all graphs represents 100% and 50% remaining mRNA expression; graphical elements created with Biorender; Fakih et. al.).

We next investigated whether T-FMRN exhibited better *in vivo* performance compared to T-DNA by measuring tissue accumulation and target silencing efficacy following subcutaneous (s.c.) administration. Wild-type adult mice (n=4/group) received a single s.c. dose (40nmol siRNA dose, equivalent to 20 mg/kg siRNA dose) of a non-targeting T-FMRN control (T-control; no antisense strands loaded), an *Htt*-targeting T-FMRN (T-3xHTT), or an HTT-targeting T-DNA (T-3xHTT-DNA) (Fig. 2c).Two weeks post-injection, mice were euthanized, and siRNA accumulation (via PNA hybridization assay) and *Htt* mRNA silencing (via Quantigene singleplex assay) were measured across major tissues to correlate structural integrity with therapeutic performance. The FMRN architecture mediated significantly higher siRNA accumulation and target silencing in the heart, lungs, and liver compared to T-DNA (Fig. 2d,e). While T-3xHTT enhanced delivery to the lung, no silencing was observed in that tissue, potentially due to the dose being lower than the silencing threshold. Notably, siRNA accumulation and target silencing in the kidney were markedly higher in the T-DNA group than in the FMRN group (∼4 vs 1 pmol/mg, ****p<0.0001 for accumulation; ∼25% vs 0%, *p<0.05, for silencing), suggesting that the unstable T-DNA scaffold is rapidly degraded and filtered into the kidneys for elimination as unconjugated siRNAs, while T-FMRN avoids rapid clearance, facilitating higher systemic retention and broader extrahepatic distribution. These results demonstrate that full chemical modification is a requirement for the biological viability and extrahepatic delivery of RNA nanostructures.

### Characterization of T- and S-FMRN shows uniform assembly with enhanced systemic pharmacokinetics *in vivo* comparable to lipophilic conjugation

We next aimed to study the impact of FMRN molecular weight (MW) and shape on siRNA pharmacokinetics and pharmacodynamics (PK/PD) *in vivo*. We assembled and characterized the MW and size of 3 architectures (Figure 3a): Arm-siRNA (unstructured, 27 KDa), T-FMRN (structured, 80 KDa), and S-FMRN (structured, 110 KDa), using gel electrophoresis and dynamic light scattering (DLS). Gel electrophoresis confirmed the uniform, monodisperse assembly of the geometries, with S-FMRN exhibiting predictably lower electrophoretic mobility compared to T-FMRN and Arm-siRNA, consistent with its higher MW (Fig. 3b). DLS analysis revealed near-identical hydrodynamic diameters for both T-FMRN (∼12 nm) and S-FMRN (∼13 nm), suggesting that despite differences in valency/MW, the overall spatial footprint of the two remains similar. We were unable to acquire a clean DLS measurement for Arm-siRNA, due to its unstructured nature.

**Figure 3:**
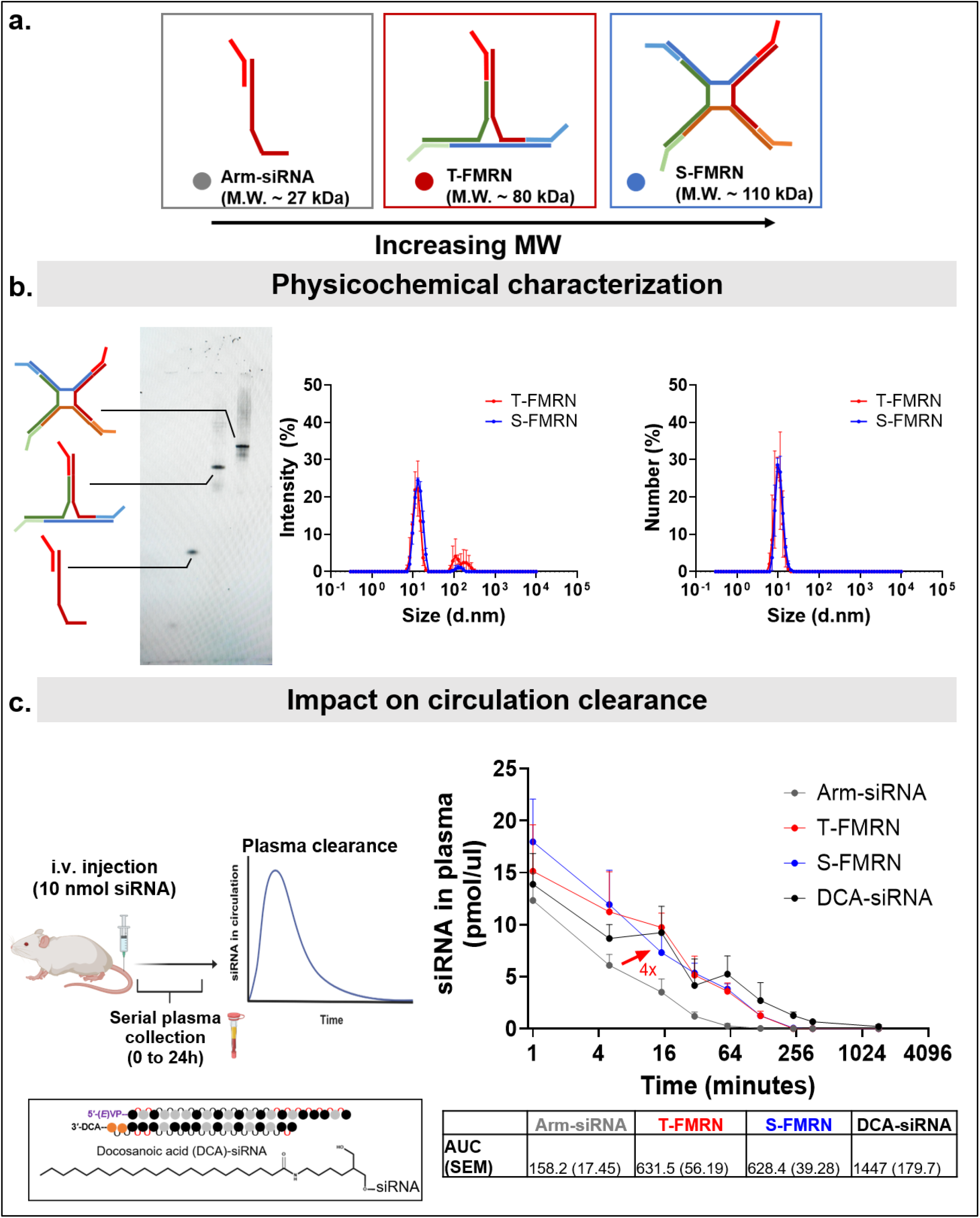
Physicochemical characterization and pharmacokinetic profiles of FMRNs. **A.** Schematic illustration of fully modified linear Arm-siRNA and polygonal T- and S-FMRN architectures. The triangular and square configurations carry 3 and 4 siRNA copies, respectively, with calculated molecular weights indicated for each assembly. **b.** Structural and solution-phase characterization. (Left) Native PAGE (8%) analysis showing distinct electrophoretic mobility shifts corresponding to the increased molecular weight of the assembled scaffolds. (Right) Hydrodynamic size distribution measured by dynamic light scattering (DLS), represented as intensity and number percentages. T- and S-FM-RNs exhibit monodisperse diameters of approximately 10 nm, whereas linear Arm-siRNA lacks a measurable monodisperse peak at this scale. **C.** *In vivo* pharmacokinetic evaluation. (Left) Schematic of the experimental timeline for intravenous (i.v.) administration (20 nmol siRNA dose, equivalent to 10 mg/kg siRNA dose). (Right) Plasma clearance kinetics quantified via serial blood collection over 24h. *Note:* plasma clearance data of DCA-siRNA is adapted from previously published work *Fakih et. al. MTNA 2025.* (Electrophoresis was performed on 8% non-denaturing PAGE gels at 130 V for 1 h in 1× TBE buffer. DLS measurements are n=2 technical replicates, with concentrations of FM-RNs at 25uM. All study groups are *n* = 3-4 mice/group. siRNA in plasma concentration was measured by PNA assay; data represented as mean ± S.D.; graphical elements created with Biorender; Fakih et. al.).

We next evaluated the impact of these nanostructures on *in vivo* PK profiles following systemic administration (Fig. 3c). We hypothesized that, unlike unstructured, unconjugated siRNA or single-arm siRNA strands, the defined nanometer-scale geometry of FMRNs would fundamentally alter clearance kinetics. Wild-type adult mice received a single intravenous (i.v.) dose for maximum bioavailability (20 nmol siRNA dose, equivalent to 10 mg/kg siRNA dose) of either unstructured Arm-siRNA, T-3xsiRNA, S-4xsiRNA, or a lipid-conjugated control (DCA-siRNA) previously validated to achieve extended plasma circulation and extrahepatic delivery.^43–45^, Serial plasma collection and siRNA quantification over 24 hours revealed that FMRN assembly significantly extended circulation time, yielding an approximate four-fold increase in the area-under-the-curve (AUC) compared to the unstructured Arm-siRNA (Fig 3c). While Arm-siRNA was rapidly cleared within ∼5 minutes, both T- and S-FMRNs exhibited an improved half-life of ∼18 minutes. This difference in clearance between the Arm-siRNA and the FMRNs further supports the intactness of the structure in circulation. Notably, the PK profiles of T- and S-FMRNs were nearly indistinguishable despite a 20 kDa difference in MW and different shape. Importantly, their circulation kinetics were comparable to those of the DCA-siRNA control (which still had a larger AUC) suggesting that clearance is likely more influenced by size and shape of FMRNs rather than the MW. To explore this further, we determined the protein binding profile of FMRNs versus DCA-siRNA following systemic administration (Supplementary Fig. 4). We found that FMRNs bind exclusively to albumin, whereas DCA-siRNA binds all present lipoproteins to various extents as previously reported.^42^ Collectively, these results identify FMRNs as highly efficient, single-material delivery systems capable of boosting tissue exposure which other studies have shown to correlate with higher tissue accumulation and therapeutic availability of siRNA payloads. This boosted siRNA circulation is comprable to those achieved with lipid conjugation strategies without any lipid moieties on FMRNs.

### FMRNs outperform siRNA cocktails in extrahepatic delivery and multi-target silencing

Following the characterization of their PK advantages, we evaluated the capacity of FMRNs to deliver multiple distinct siRNAs simultaneously. We engineered T-FMRNs carrying three distinct siRNAs, each targeting *Htt, Jak1*, or *Mecp2* mRNAs (T-3xTargets) and S-FMRNs carrying four different siRNAs targeting *Htt*, *Jak1*, *Mecp2*, or Msh3 mRNAs (S-4xTargets) (Fig. 4a). T-3xTargets and S-4xTargets FMRNs were administered s.c. at a dose of 40 nmol of scaffold (equivalent to 40 nmol/siRNA target, corresponding to 20 mg/kg per siRNA target; detailed dosing calculations in Supplementary Fig. 5a). As a comparative control, we used a “cocktail” of four unconjugated, fully chemically stabilized siRNAs (UNC-4xTargets), targeting the same four genes as S-FMRNs. UNC-4xTargets was dosed at 40 nmol/siRNA target (equivalent to 20 mg/kg/target, and a total of 160 nmol of scaffold) — matching the per-target molar siRNA dose of the FMRN but delivering four times the molar scaffold dose. Efficacy was compared to a non-targeting T-FMRN control (scaffold structure not loaded with antisense strands).

**Figure 4:**
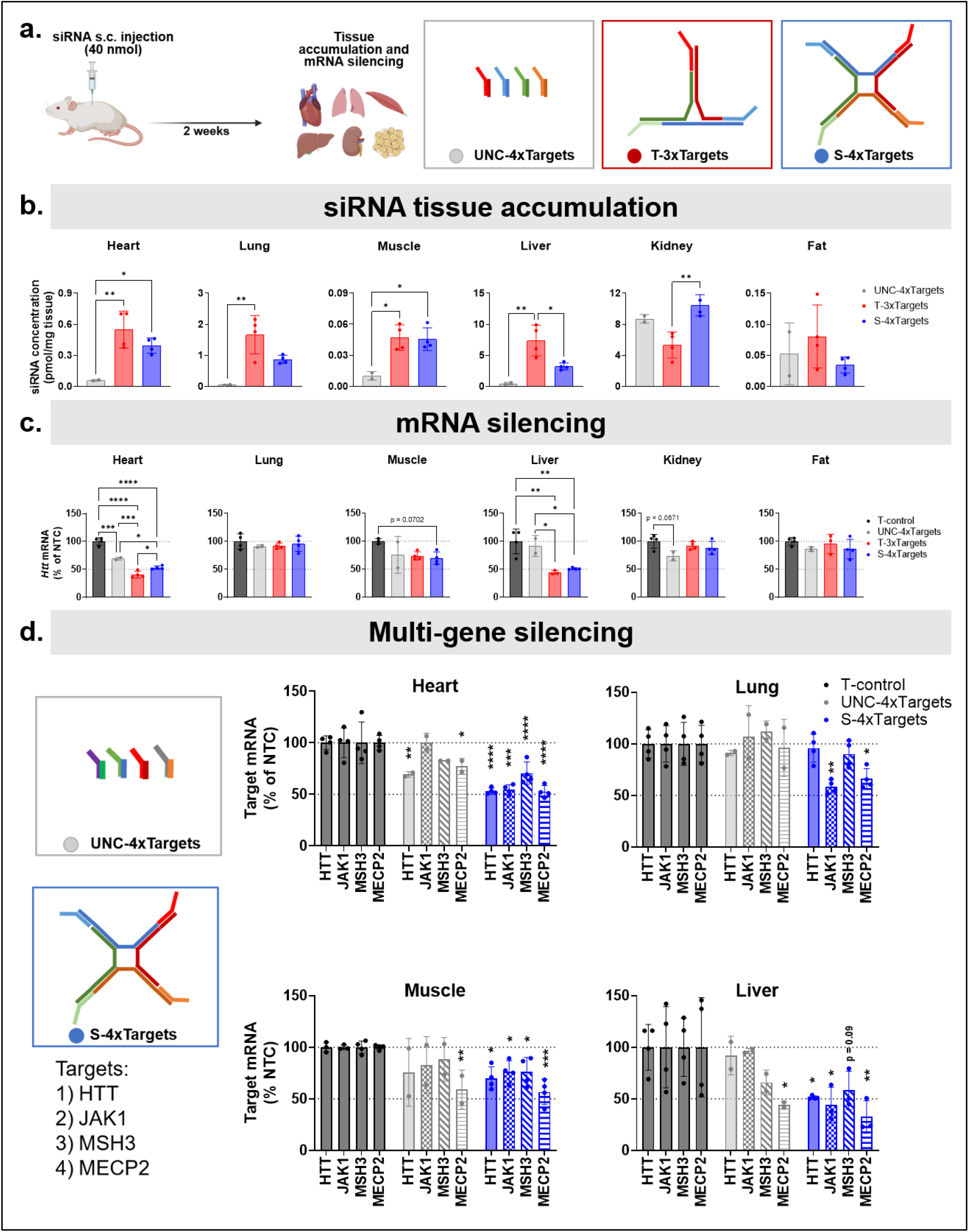
T- and S-FMRN improve siRNA tissue accumulation, target mRNA silencing, and multi-gene targeting in various extra-hepatic tissues following systemic injection *in vivo*. **a.** Scheme of *in vivo* study evaluating the distribution and efficacy of integrated multi-gene targeting. Mice received subcutaneous (s.c.) injections of either an unconjugated siRNA cocktail (UNC-4xTargets), T-3xTargets, or S-4xTargets (40 nmol siRNA dose/target, equivalent to 20 mg/kg siRNA dose/target). Tissues were harvested two weeks post subcutaneous injection to assess siRNA accumulation and mRNA silencing of HTT, JAK1, MSH3, and MECP2 mRNAs. **b.** siRNA Tissue accumulation and **c.** HTT mRNA silencing profiles across major extrahepatic tissues. Efficacy was benchmarked against a non-targeting control (T-FMRN lacking therapeutic antisense strands). **d** Simultaneous multi-gene target silencing. mRNA expression levels for HTT, JAK1, MSH3, and MECP2 were compared between the unconjugated UNC-4xTargets cocktail and the integrated S-4xTargets architecture in tissues exhibiting high accumulation levels. All study groups are *n* = 3-4 mice/group. Statistical analysis was performed in GraphPad Prism: for c. one-way ANOVA analysis followed by Tukey’s multiple comparisons across all groups for HTT silencing; for d. two-way ANOVA analysis followed by Dunnet’s multiple comparisons comparing to T-control for every target mRNA expression; non significant comparisons not shown, \**p* < .05, \*\**p* < .01, \*\*\**p* < .001, \*\*\*\**p* < .0001. (tissue accumulation measured by PNA assay; mRNA measured by the Quantigene 2.0 Assay; data represented as mean ± S.D.; dotted line on all graphs represents 100% and 50% remaining mRNA expression; graphical elements created with Biorender; Fakih et. al.).

Two weeks post-injection, we quantified accumulation of the *Htt*-targeting siRNA to benchmark distribution across modalities (Fig. 4b). Despite receiving only 25% of the total scaffold dose compared to the unconjugated siRNA cocktail, FMRNs mediated significantly higher siRNA accumulation across all extrahepatic tissues analyzed, including heart (∼5-fold), lung (10-fold), muscle (3-fold), and liver (3-fold). Intriguingly, in the liver, the triangular scaffold supported significantly higher siRNA accumulation than the square scaffold (7 vs. 4 pmol/mg; *p < 0.05), despite the square scaffold’s higher MW. Conversely, the square scaffold supported higher renal accumulation of siRNA (8 vs. 4 pmol/mg, **p < 0.01), potentially reflecting geometry-dependent filtration kinetics. Calculating percentage of injected dose (%ID) that accumulates in these tissues (Supplementary Fig. 5b), we again observe that FMRNs strongly enhances extrahepatic delivery with higher %ID compared to the mixtures of UNC-siRNAs in all measured tissues except for kidney where most of unconjugated naked siRNAs end up in.

The superior biodistribution of FMRNs directly translated into potent, concurrent multi-gene silencing. Evaluating *Htt* mRNA expression as the primary outcome (Fig. 4c), both the triangular and square scaffold groups achieved significant silencing in the heart (∼50% knockdown, ****p < 0.001) and liver (∼50% knockdown, ****p < 0.001). The triangle scaffold exhibited a slight, but significantly higher silencing effect in the heart compared to square scaffold (60% vs. 50% of control, *p < 0.05). The square scaffold also mediated near-significant silencing in skeletal muscle (∼20% knockdown, p = 0.07). In contrast, the unconjugated siRNA cocktail achieved only modest silencing in the heart (∼25%, ***p < 0.01)—significantly less than the FMRN architectures (*p < 0.05)—and failed to achieve any significant silencing in the liver (*p < 0.05 compared to FMRNs).

We next assessed simultaneous silencing across all targets in organs with the highest accumulation differential to the control (Fig. 4d). In the heart, the square scaffold achieved robust (∼50%) silencing across all four targets (****p < 0.001), whereas the unconjugated siRNA cocktail exhibited only modest modulation of *Htt* and *Mecp2* mRNAs (∼20%, *p < 0.05). In the liver, the square scaffold facilitated concurrent silencing of all four targets (∼50% knockdown; *p < 0.05), whereas the cocktail showed significant modulation of *Mecp2* mRNA only. Within the lung, the square scaffold achieved significant silencing of *Jak1* mRNA (∼50% knockdown, **p < 0.01) and *Mecp2* mRNA (∼30% knockdown, *p < 0.05), but did not affect *Htt* or *Msh3* mRNAs expression, suggesting that geometry-specific cellular uptake or intracellular trafficking could dictate efficacy in specific pulmonary cell types. Finally, in skeletal muscle, the square scaffold achieved modest but concurrent silencing of all four targets (ranging from ∼20% knockdown for *Htt*, *Jak1*, and *Msh3* mRNAs, *p < 0.05, to ∼50% for *Mecp2* mRNA, ***p < 0.001), whereas the cocktail modulated only *Mecp2* mRNA to similar levels as the square scaffold (∼40% knockdown, ** p < 0.01). Collectively, these data demonstrate that integrating multiple siRNAs into a single, structurally defined FMRN entity provides an advantage over therapeutic unconjugated siRNA mixtures, enabling synergistic multi-gene modulation with enhanced potency and molar efficiency.

In a follow up study, we also evaluated the durability of these effects by assessing tissue accumulation and silencing efficacy at one month following a single s.c. injection, where we further included the lipophilic DCA-conjugated siRNA as a benchmark modality that supports extrahepatic siRNA delivery and currently in preclinical and clinical development (Supplementary Fig. 6).^13,15,46,47^ Both T-FMRN and S-FMRN maintained siRNA tissue accumulation and target gene silencing levels comparable to those of DCA-siRNA across all evaluated tissues (heart, lung, skeletal muscle, and liver),with DCA-siRNA showing stronger silencing only in the muscle (∼50% vs ∼20%, for DCA vs T and S-FMRNs, ***p<0.001). FMRNs also had overall comprable %ID to that of DCA-siRNA in the extrahepatic tissues assessed (Supplementary Fig. 7), with FMRNs having higher %ID in kidney compared to DCA (∼2% for T- and S-FMRNs vs 1% for DCA-siRNA, ***p<0.001), and both DCA-siRNA and S-FMRN having significantly higher accumulation in fat compared to T-FMRN (0.2% for DCA and S-FMRN, vs 0.08 for T-FMRN, *p<0.05). Overall, these data further supports the durability of effects achieved with FMRNs, to similar levels as those observed with standard delivery technologies like lipophilic conjugates acting as a benchmark modality.

We also assessed FMRN-mediated silencing at the protein level by targeting myostatin (Mstn) in muscle, which allows for longitudinal monitoring of efficacy since MSTN is secreted into plasma (Supplementary Fig. 8). Using a previously validated system for continuous MSTN monitoring^48^, mice were injected s.c. with T-FMRN (40 nmol siRNA dose) targeting Mstn, and plasma was collected weekly for four weeks to measure MSTN protein levels, alongside terminal Mstn mRNA measurement in muscle tissue at the four-week endpoint. T-FMRN supported approximately 50% silencing of plasma MSTN protein from week one through week three, with slight recovery observed at week four (∼40% silencing). This protein-level silencing was consistent with Mstn mRNA silencing measured at the four-week endpoint, where approximately 50% mRNA silencing was still observed.

Lastly, we validated that the FMRN scaffold oligonucleotides are not contributing to any biological effects of efficacy by generating a T-FMRN that carries 3 non-active NTC sequences (T-3xNTC). We compared its ability to silence JAK1 in a dose-dependent study *in vitro* to that of T-FMRN and S-FMRN that do carry an siRNA against JAK1 (T-3xTargets and S-4xTargets), and found that indeed T-3xNTC does not induce any JAK1 silencing (Supplementary Fig. 9). This supports the notion that the scaffold oligonucleotides are not contributing to RNA interference.

### FMRNs exhibit a favorable systemic safety profile

To complement the PK/PD results, we evaluated the safety of FMRNs following s.c. injection. Specifically, we recorded body weights pre injection and over a period of two weeks, as well as assessed hematological and biochemical parameters longetutidenally at 1,3, and 14 days post post-injection to probe any acute and chronic pertubations. We compared FMRN-treated mice (T-3xTargets, S-4xTargets) with PBS control and the mixture of unconjugated siRNAs (UNC-4xsiRNAs). All groups were dosed at 40 nmol siRNA/target (equivalent to 20 mg/kg) as in the previous study (Supplementary Fig. 10). No significant changes to body weights were observed (Supplementary Fig. 10a). Quantitiatve analysis of complete blood counts (CBC) and comprehensive metabolic chemistry revealed that FMRNs were well-tolerated, with no evidence of systemic toxicity (Supplementary Fig 10 and 11). Markers of hepatotoxicity (e.g., ALT) and nephrotoxicity (e.g., BUN) remained within normal physiological ranges across all treatment groups (Supplementary Fig. 10b), and total platelet counts and immune cell populations (including neutrophils, lymphocytes, and monocytes) exhibited no significant deviations from baseline, supporting the safety of the platform. There was a significant slight increase in ALT and percentage of neutrophils at 1-day post injection with the FMRNs, but the increase was still within normal ranges for these readouts, and the increase faded away by day 3 and not observed at day 14. The overall data supports the safety of FMRN and the absence of acute or chronic hematological stress or immunogenic reactions. Similarly, we did a measurement at day-3 post injection of the study comparing to DCA-siRNA (Supplementary Fig. 6 and 12) and linear Arm-siRNA (non-structured control), with similar results obatained of no evidence of systemic toxicity. Collectively, these data support a favorable safety profile for FMRNs at the tested doses and time points.

### FMRNs support safe, effective, and durable multi-gene silencing in the CNS

To evaluate the broader applicability of the FMRN platform, we investigated its capacity for local delivery gene modulation within the CNS, a site of significant therapeutic interest for polygenic neurodegenerative disorders such as Huntington’s disease (HD) and Alzheimer’s disease (AD). We loaded T-FMRNs with three distinct, previously validated siRNAs (T-3xTargets) targeting *Htt, Jak1,* or *Msh3*—genes implicated in HD and AD. Wild-type adult mice received bilateral intracerebroventricular (ICV) injections of either T-3xTargets, a clinical-stage *Htt*-targeting divalent siRNA scaffold (Divalent-HTT) as a positive control ^8,49^, a standard unconjugated siRNA targeting *Htt* (UNC-HTT) or a non-targeting control (UNC-NTC). All doses were at 10 nmol of scaffold (10 equivalent to 10 nmol/siRNA target) (Fig. 5a).

**Figure 5:**
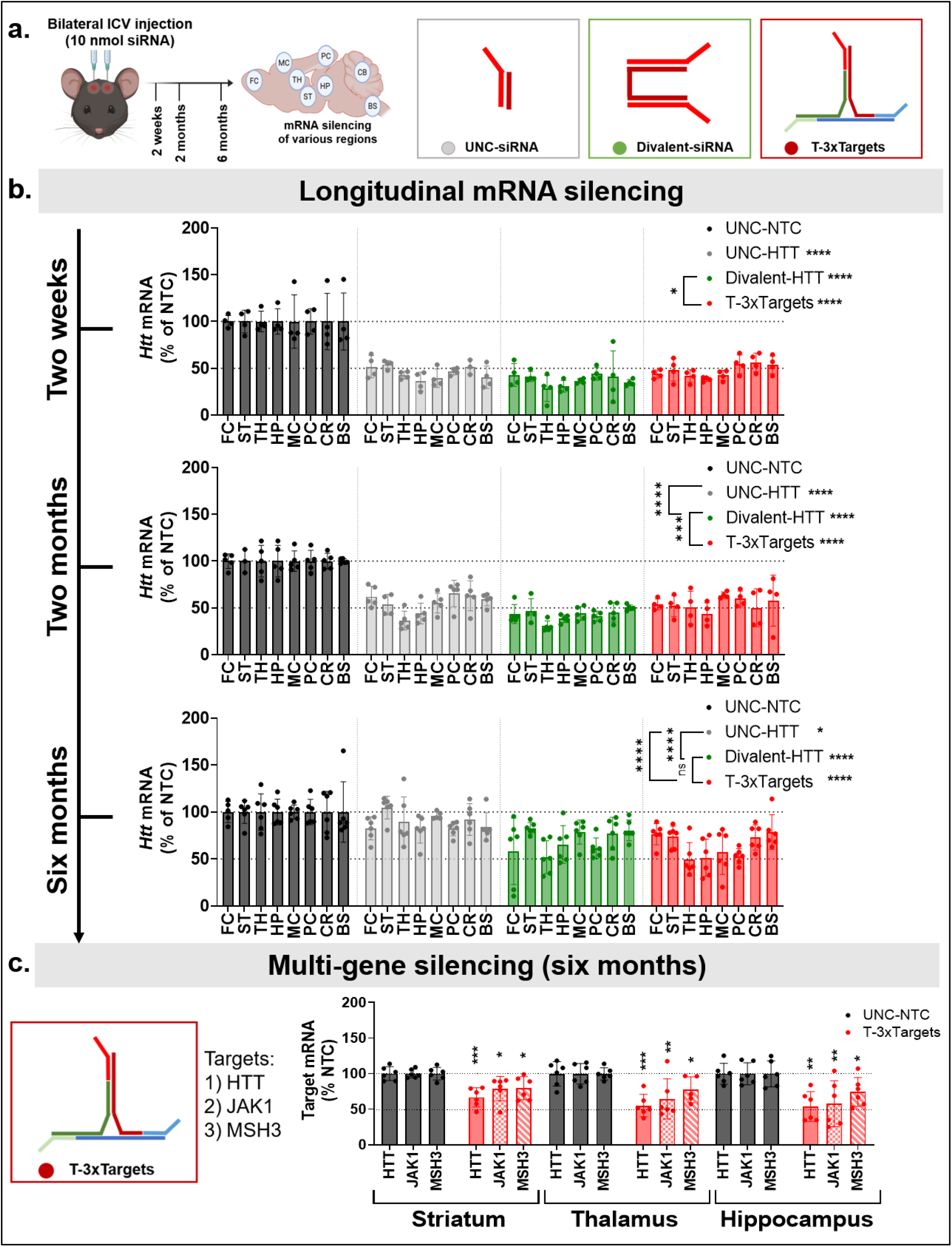
T-FM-RN facilitate potent and long-term multi-gene silencing in the central nervous system. **a.** Scheme of *in vivo* study, where mice were intracerebroventricularly (ICV) injected with HTT-targeting UNC-siRNA, Divalent-siRNA, or T-FM-RN targeting 3 targets (T-3xTargets), at a 10 nmol siRNA dose (10nmol HTT-siRNA for all groups, 10nmol/target for T-FMRN, equivalent to 10 nmol scaffold dose). mRNA silencing in various brain regions was measured at two-weeks, two-months, and six-months post injection. (brain regions: Frontal cortex-FC; Striatum-ST; Thalamus-TH; Hippocampus-HP; Medial cortex-MC; Posterior cortex-PC; Cerebellum-CS; Brain stem-BS). **b.** Longitudinal HTT mRNA silencing at two-weeks, two- and six-months post injection, with siRNA activity compared to non-targeting control UNC-NTC. **c.** Simultaneous multi-gene target silencing (HTT, JAK1, MSH3 mRNA) with T-3xTargets in striatum, thalamus and hippocampus, at six-months post ICV injection, compared to UNC-NTC control. All study groups are *n* = 4-6 mice/group. Statistical analysis was performed in GraphPad Prism: for b. two-way ANOVA analysis across all groups; for c. two-way ANOVA analysis followed by Dunnet’s multiple comparisons comparing to control for every target mRNA expression; \**p* < .05, \*\**p* < .01, \*\*\**p* < .001, \*\*\*\**p* < .0001. (mRNA measured by the Quantigene 2.0 Assay; data represented as mean ± S.D.; dotted line on all graphs represents 100% and 50% remaining mRNA expression; graphical elements created with Biorender; Fakih et. al.).

*Htt* mRNA expression was assessed across brain regions at two-weeks, two-months, and six-months post-injection to determine both silencing efficacy and duration of effect (Fig. 5b). At the two-week timepoint, all groups resulted in robust HTT mRNA silencing across all regions (***p<0.0001 vs UNC-NTC). Divalent-HTT showed a slight but significantly higher efficacy compared to T-3xTargets (*p<0.05), despite siRNA accumulation across all regions trending higher in the T-3xTargets group compared to UNC-HTT and Divalent-HTT (Supplementary Fig. 13a). Two-months post injection, T-3xTargets maintained robust and widespread silencing of HTT mRNA across various brain regions (****p < 0.0001 vs. UNC-NTC), with the Divalent-HTT still exhibiting slight but significantly higher efficacy (***p < 0.001 vs T-3xTargets), particularly in posterior regions such as the cortex, cerebellum, and brainstem (Fig. 5b). At six-months post injection, the silencing efficacy of T-3xTargets was comparable to that of Divalent-HTT scaffold, with both significantly outperforming the unconjugated UNC-HTT siRNA (****p<0.0001 vs UNC-HTT for both groups), which lost nearly all efficacy across most brain regions (Fig. 5b).

Importantly, mice injected with FMRNs exhibited no apparent toxicity, with 100% survival to the designated endpoints, supporting a favorable safety profile for direct CNS administration. This was further validated by quantifying mRNA levels of *Iba1* (microgliosis marker) and *Gfap* (astrogliosis marker) two months post-injection in the thalamus and hippocampus—the regions exhibiting the strongest silencing and highest accumulation, and closest to the siRNA injection site. No significant changes in these inflammatory markers were observed across any of the treatment groups, supporting the safety profile of FMRNs in the CNS, by not eliciting an immune response or neurotoxicity at therapeutic doses used in this study (Supplementary Fig. 13b).

To evaluate the multi-gene targeting capacity of the FMRN platform in the CNS, we measured the expression of all three targets (*Htt*, *Jak1*, and *Msh3* mRNAs) in the striatum, thalamus, and hippocampus (highest observed accumulation and activity, being closest to site of injection) six months post-injection. T-3xTargets successfully maintained silencing of all three genes across these regions (ranging from 15-50% for various targets in various regions, *p<0.05 to ***p<0.001) (Fig. 5c). These results demonstrate that FMRNs offer efficacious pharmacological durability in the CNS, providing a versatile and long-lasting treatment strategy for complex neurological conditions.

### FMRN modularity reveals MW dependencies for siRNA activity in the brain

Following validation of the safety and long-term efficacy of FMRNs in the CNS, we leveraged the platform’s modularity to investigate the fundamental factors governing siRNA distribution in the brain. While previous studies have suggested that increased MW or size may correlate with broader CNS distribution—as exemplified by divalent siRNA scaffolds^8^—the individual contribution of these parameters remains poorly defined. FMRNs provide a unique opportunity to decouple these factors, as T- and S-FMRNs have different MWs despite possessing similar hydrodynamic diameters.

We systematically interrogated the impact of MW on brain efficacy by administering a series of scaffolds with increasing MW (UNC-siRNA, Divalent-siRNA targeting HTT, Arm-siRNA targeting HTT, T-3xTargets (targeting *Htt*, *Jak1*, and *Mecp2* mRNAs), and S-4xTargets (targeting *Htt, Jak1, Mecp2,* and *Msh3* mRNAs) via ICV injection at a total dose of 5 nmol HTT-siRNA (Fig. 6a). At one month post-injection, all architectures achieved statistically significant *Htt* mRNA silencing across multiple brain regions (Fig. 6b) (****p<0.0001 all groups vs UNC-NTC). Intriguingly, we observed distinct MW-dependent trends in silencing efficacy, most notably at the front and back brain. In the frontal cortex, we observed a direct correlation between MW and potency, where S-4xTargets was the sole scaffold to achieve significant silencing (∼15%, *p<0.05) (Fig. 6c). Conversely, in the brain stem, increasing MW led to a steady significant decrease in efficacy. Here, the lower-MW UNC-siRNA mediated the strongest silencing effects (∼45% silencing, ****p<0.0001 vs UNC-NTC), whereas efficacy dropped as we went up in MW: 35% silencing for Divalent-siRNA, 19% for Arm-siRNA, 13% T-3xTargets and 4% for S-4xTargets (Fig. 6c). These findings suggest that although molecular size may influence initial CSF clearance, the chemical architecture, molecular weight, and valency of a compound play important roles in determining regional tissue accumulation, residence time, and potency, highlighting the need for further systematic evaluation.

**Figure 6:**
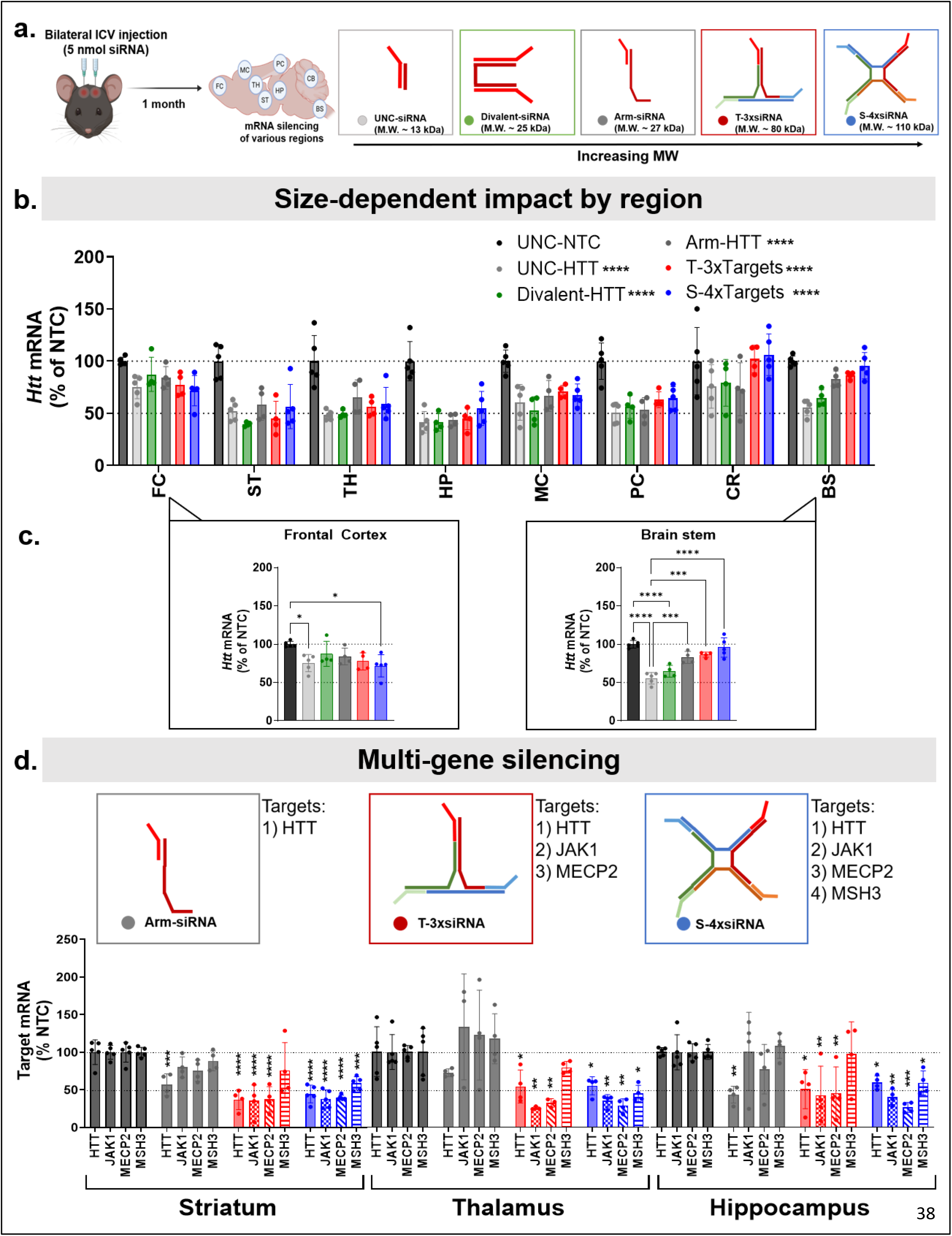
Systematic interrogation of molecular weight-dependent regional distribution in the brain. **a.** Scheme of *in vivo* study for decoupling molecular weight and hydrodynamic size. Mice were injected via ICV with a fixed 5 nmol HTT-targeting siRNA dose (equivalent to 5 nmol scaffold dose for all) across a series of scaffolds with increasing molecular weight: UNC-siRNA, Divalent-siRNA, Arm-siRNA, T-3xsiRNA, and S-4xsiRNA. mRNA silencing in various brain regions was measured (brain regions: Frontal cortex-FC; Striatum-ST; Thalamus-TH; Hippocampus-HP; Medial cortex-MC; Posterior cortex-PC; Cerebellum-CS; Brain stem-BS). **b.** HTT mRNA silencing at one-month post injection, with siRNA activity compared to non-targeting control UNC-NTC. **c.** Weight-dependent efficacy trends in the front(frontal cortext) and back (brain stem) of the brain. **d.** Simultaneous multi-gene target silencing (HTT, JAK1,MECP2, MSH3 mRNA) with T-3xTargets and S-4xTargets in striatum, thalamus and hippocampus, compared to UNC-NTC control. All study groups are *n* = 3-4 mice/group. Statistical analysis was performed in GraphPad Prism: for b. two-way ANOVA analysis across all groups; for c. one-way ANOVA analysis followed by Tukey’s multiple comparisons across all groups; for d. two-way ANOVA analysis followed by Dunnet’s multiple comparisons comparing to control for every target mRNA expression; non significant comparisons not shown, \**p* < .05, \*\**p* < .01, \*\*\**p* < .001, \*\*\*\**p* < .0001. (mRNA measured by the Quantigene 2.0 Assay; data represented as mean ± S.D.; dotted line on all graphs represents 100% and 50% remaining mRNA expression; graphical elements created with Biorender; Fakih et. al.).

Finally, we validated the platform’s multi-gene silencing capabilities following ICV administration (Fig. 6d). Target mRNA silencing for Arm-siRNA (targeting *Htt* only), T-3xTargets (targeting *Htt, Jak1, Mecp2*), and S-4xTargets(targeting *Htt, Jak1, Mecp2, Msh3*) was evaluated in the striatum, thalamus, and hippocampus as previously done. While Arm-siRNA silenced *Htt* mRNA alone, T-3xTargets successfully silenced all three targets across all tested regions without affecting the non-targeted *Msh3* mRNA. Importantly, the S-4xTargets architecture achieved potent (∼50%) and simultaneous silencing of all four genes across all regions and targets analyzed (Fig. 6d). These results demonstrate the versatility of the FMRN platform for potent multi-gene silencing, with precise control over the number of targets modulated.

## DISCUSSION

The development of fully chemically-modified RNA nanostructures (FMRNs) represents a promising convergence of structural nanotechnology and oligonucleotide therapeutics. While nucleic acid nanotechnology has achieved remarkable architectural and design advances, effective *in vivo* translation for drug delivery has been limited by insufficient metabolic instability and structural complexity. Here we demonstrate that oligonucleotides fully stabilized with chemical modifications that are clinically validated (2ʹ-F and 2ʹ-OMe, phosphorothioate linkages and 5’-vinyl phosphonate) can assemble into functional nanostructures that retain therapeutic RNAi efficacy (compatible with Ago2 loading). To our knowledge, this is the first report of fully chemically modified, vehicle-free RNA nanostructures capable of effective and sustained extrahepatic delivery and multi-gene silencing, establishing a new direction at the interface of RNA nanotechnology and therapeutic oligonucleotide design.

Substantial evidence from preclinical and clinical studies indicates that manipulation of biophysical properties improves oligonucleotide pharmacokinetics and tissue exposure.^3^ For example, divalent and tetravalent siRNA architectures have demonstrated enhanced retention and efficacy in the CNS, eye, and lung^8,50,51^, while lipophilic conjugates have similarly improved tissue accumulation and clinical utility.^45,52,53^ In each case, therapeutic performance is improved through modulation of molecular size, valency, or physicochemical properties. The FMRNs here extend this principle by demonstrating that fully modified oligonucleotides can be assembled into larger, structurally defined entities without requiring additional carrier materials. Notably, the increased thermodynamic stability of fully modified oligonucleotides facilitates nanostructure assembly, simplifying design relative to conventional DNA-based nanostructures and expanding the accessible structural space.

The resulting nanostructures exhibit altered biophysical properties that enhance circulation time, tissue accumulation, and functional delivery following both local and systemic administration. Importantly, these pharmacokinetic advantages are achieved without the addition of nonfunctional carrier components. This feature distinguishes FMRNs from many delivery technologies, including antibody-oligonucleotide conjugates, where the vehicle contributes the majority of the molecular mass. In contrast, the majority of the mass within FMRNs consists of therapeutically active oligonucleotide sequences, enabling simultaneous manipulation of pharmacokinetic properties while preserving high active-drug content. This maximizing of active-drug content enables the integration of multiple distinct siRNAs within a single framework to address multigenic diseases. Furthermore, as a single-material system, the vehicle-free FMRN circumvent some of the manufacturing complexity, batch-to-batch variability, toxicity, and solubility challenges inherent to multi-component formulations (e.g, LNPs).

A key conclusion from this study is that complete chemical stabilization appears to be a fundamental design requirement, not just an optimization strategy, for successful *in vivo* application of nucleic caid nanostructures. While partial modification strategies (e.g., localized phosphorothioate incorporation and stabilization of vulnerable regions) can provide measurable protection *in vitro*, the *in vivo* environment presents a substantially greater challenge.In biological tissues, oligonucleotides are exposed to a diverse and highly active repertoire of nucleases that rapidly exploits points of metabolic vulnerability, which inititate degradation and ultimately disassembling the nanostructure. Consequently, stabilization strategies that appear effective in serum or cell culture often fail to provide the durability required for sustained *in vivo* function. This finding mirrors therapeutic siRNA studies by Hassler et al., where partially modified siRNAs lacked the *in vivo* durability of fully stabilized counterparts^39^, emphasizing the importance of eliminating isolated points of metabolic vulnerability. Our findings extend this principle to higher-order architectures, as evidenced by the dramatic difference in stability and efficacy between FMRNs and partially stabilized DNA structures, with FMRNs achieving prolonged pharmacological activity, including six-month durability in the CNS.

Critically, the FMRN platform provides a unique architecture for multi-dimensional, combinatorial therapeutic delivery with precision, within a singular molecular framework. ^54,55^ While traditional carriers rely on random physical encapsulation^55^ and multi-drug cocktails suffer from uncoordinated *in vivo* pharmacokinetics, a single-entity FMRN with precise structural control ensures synchronized cellular uptake and reproducible, coordinated silencing across all targets.This enables the co-delivery of distinct modalities (e.g., ASOs and siRNAs) using “rational valency”, such as a square-FMRN engineered with a 3:1 copy ratio to match a disease’s differential gene expression and allowing stochiometric weighting. Overall, this single chemistry entity has the potential to streamline future translational and regulatory approval pipeline, eliminating the exhaustive, multi-arm safety and efficacy trials required for combination therapies.

Additionally, the modularity of the FMRN allowed us to decouple the effects of MW and hydrodynamic size on CNS distribution. The inverse efficacy trends observed between the frontal cortex and brain stem as molecular weight increased reveal a size-dependent regional partitioning, offering a new programmatic handle to tune architectures for specific anatomical focal points in the brain.

Despite these capabilities, several limitations remain to be explored. First, future efforts must evaluate FMRN performance against broader benchmarks, such as PEGylated systems. Second, a systematic evaluation of the scaffold modification patterns is a necessary exploration as it could yield more compact or potent architectures. Third, while vehicle-free FMRNs were comparable to lipophilic conjugates systems (DCA-siRNA), incorporating lipophilic character remains highly valuable for potentially maximizing absolute tissue retention, cellular uptake, and endosomal escape. This however could be incorporated, by utilizing the modularity of the FMRN platform that allows for functionalization with lipid-or conjugate-based systems to create a synergistic framework. Recent evidence suggests that decorating nanostructures with lipids can control the polarity and hydrophobicity of the scaffold, potentially enabling cell-type-specific targeting or enhanced endosomal escape.^56^ This also highlights the need for more comprehensive investigation of the cellular uptake and endosomal escape pathways of this platform in future studies. Lastly, while the similar size but different molecular weight between T-and S-FMRN allowed to us to decouple the effects of these parameters, this remains a limited subset that requires further systematic investigation, such as varying size within the same shape, and vice-versa, which will be the focus of future work. Overall, such chemical and geometric tuning will be essential to minimize systemic clearance, with the goal of maximizing bioavailability via physicochemical control.

In summary, this work moves the field of nucleic acid nanotechnology a step closer towards robust *in vivo* drug delivery applications, particularly in the contexts of multi-gene targeting and vehicle-free control of delivery and pharmacokinetics. By establishing that fully chemical stabilization is critical, we have defined a technology that circumvents historical stability barriers. Future investigations into the interplay between geometry, core chemistry, and functionalization of the programmable FMRNs will be essential to fully realize the potential of the platform.

## METHODS

### Design of polygon arms for the assembly of RNA nanostructures

Sequences for the assembly of the triangle and square RNA nanostructures were manually designed and validated for secondary interactions, with a starting base design based on work by McLaughlin et al.^57^ Sequences designed to be complementary to each other in the scaffold were manually validated to not have any secondary interaction with other sequences in the scaffold for more than 5 complementary nucleotides. Triangle nanostructures, regardless of the variant, were assembled from polygon arms termed T1, T2, and T3, and Square nanostructures were assembled from polygon arms termed T1, T2, S3, and S4 (T1 and T2 are shared between Triangle and Square). siRNA sequences targeting *JAK1/Jak1* were previously validated by Tang *et. al.* and Tang, Gross, Fakih *et. al.,* ^13,58^; *MECP2/Mecp2* targeting sequences were previously validated by Khvorova and Hariharan ^59^; *HTT*/*Htt* targeting sequences previously validated by Sarli *et. al.,* ^60,61^; and MSH3 targeting sequences were acquired from the work of O’Rielly *et. al*.^62^ All sequences and their modifications are provided in Supplementary Table 1.

### Synthesis and purification of oligonucleotides

All strands were synthesized using a MerMade 12 (BioAutomation) synthesizer. Standard RNA 2′-O-methyl and 2′-fluoro modifications were applied to improve stability (ChemGenes and Hongene). All strands were synthesized on controlled pore glass (CPG) functionalized with a Unylinker (Hongene) (500A-1000A), and the antisense strand of the siRNA was modified with custom 5′-(*E*)-vinylphosphonate 2′-OMe-uridine CED phosphoramidite (Hongene) to introduce 5′-(*E*)-vinylphosphonate at the 5ʹ end.

All strands were cleaved and deprotected using 28% aqueous ammonium hydroxide solution supplemented with 3% diethylamine, for 16 hours at 55°C, followed by drying under vacuum at 40°C and resuspension in Millipore H_2_O. Oligonucleotides were purified using an Agilent 1290 Infinity II HPLC (Agilent Technologies) on a C18 column for lipid-conjugated sense strands and a source 15Q ion-exchange column for the other strands. Purified oligonucleotides were desalted by size-exclusion chromatography and characterized by LC-MS analysis on an Agilent 6530 accurate-mass quadrupole time-of-flight (Q-TOF) LC/MS (Agilent Technologies).

### Duplex formation and structure assembly

Equimolar amounts of strands were prepared in 1x PBS and incubated at 95°C for 5 minutes, then rapidly cooled to room temperature. To validate efficient duplex formation, 2 pmol total RNA of any duplex or assembly was loaded onto a non-denaturing 8% polyacrylamide gel (PAGE) in tris-borate-EDTA (TBE) buffer (Invitrogen #EC62155BOX) and run at constant 130V for 1h for 8% PAGE. The gel was then washed stained with 1X SYBR Gold Nucleic Acid Gel Stain (Invitrogen # S11494). Bands were visualized on the ChemiDoc MP Imaging system (Bio-Rad # 17001402).

### Cell culture and dose-dependent *in vitro* studies

HeLa cells (ATCC; #CCL-2) were maintained in Dulbecco’s Modified Eagle’s Media (Corning Cellgro; #10-013CV. Media were supplemented with 10% fetal bovine serum (Gibco; #26140) and 100 U/ml penicillin– streptomycin to make complete growth media, and all cells were grown at 37 °C with 5% of CO_2_ supply. Cells were split every 2–4 days and discarded after the passage number reached 20-30. Cell counting was performed by trypan blue staining using a Cellometer Auto 1000 instrument (Nexcelom).

For seven-point dose-response studies, RNA nanostructures or siRNAs were transfected using Lipofectamine™ RNAiMAX Transfection Reagent (Invitrogen, #13778150) according to the manufacturer’s protocol. Briefly, siRNAs and RNA nanostructures were diluted in OptiMeM (Carlsbad, CA; 31985-088) to an initial concentration of quadruple their final concentration, with a 10-fold series dilution prepared. These were then mixed with RNAimax transfection reagent solution in a 1:1 ratio, resulting in twice the desired final siRNA concentration. Finally, 50 μL of this dilution was placed in triplicate on a 96-well plate, followed by the addition of 50 μL, resulting in the desired final siRNA concentration indicated in the plots. Cells were trypsinized, centrifuged, resuspended in complete growth medium, counted, and diluted to 8,000–12,000 cells per 50 μL. These cells were then added to the test groups in the 96-well plate and incubated for 72 h at 5% CO_2_ and 37°C. Final conditions are of 5% FBS supplemented growth media, 10 nM top concentration of active siRNA, followed by 6 concentrations at a 10-fold dilution, all in triplicates. mRNA levels were then quantified as described in the next section.

### mRNA quantification

mRNA expression was quantified with the QuantiGene Singleplex Assay Kit (Invitrogen # QS0016) as previously described. ^13,46^ Cultured cells were lysed in a total volume of 250 µL diluted lysis mixture consisting of one part lysis mixture, two parts water, and 0.167 mg/mL proteinase K (Invitrogen # AM2548). Lysates were then thoroughly mixed by pipetting. Probe set validation was performed for each target to determine an appropriate lysate volume that enabled unambiguous, linear mRNA detection. The corresponding volumes were transferred to the capture plate as indicated in the manufacturer’s protocol (Invitrogen # MAN0018628), and the protocol was followed. Luminescence detection was performed using the SpectroMax M5 Microplate Reader (Avantor). The following QuantiGene Singleplex RNA probes from ThermoFisher Scientific were used for mRNA detection: HTT (Human, SA-50339) and HPRT (Human, SB-15463), JAK1 (Human, #SA-50455), MECP2 (Human, SA-14665), MSH3 (Human, SA-3002246), Htt (Murine, SB-14150), Hprt (Murine, SB-15463), Jak1 (Murine, SB-3029714), Mecp2 (Murine, SB-11697), Msh3 (Murine, SB-3030208), Gfap (Murine, SB-14051), Iba-1 (Murine, SB-3027744).

For mouse tissue lysate preparation and mRNA quantification, tissues were collected at the indicated time points and stored in RNA later solution (Sigma-Aldrich; #R0901) at 4 °C overnight. Punch biopsies from the specified tissues were collected (1x4mm punches) and added to 400 μL of homogenizing solution (Invitrogen; QS0517) containing 0.2 mg/mL proteinase K (Invitrogen, #AM3546), in a QIAGEN collection microtube holding a 3-mm tungsten bead. The tissues were then homogenized for 10 min at 30 Hz using a QIAGEN TissueLyser II, followed by incubating at 55 °C for 30 min. Lysate was then used as described above for mRNA quantification, or for siRNA accumulation (described below), or stored at -20 °C.

### Serum stability assay

RNA nanostructures were prepared at 200 µM in 1xPBS as described above. Structures were then added to DMEM media supplemented with 10% FBS (the same growth media for cell culture) to dilute structures to a final concentration of 5 µM. Aliquots (10 μl, equivalent to 50 pmol) were collected at various timepoints (pre-media addition, 0 mins, 1h, 1 day, 3 days, 5 days, and 7 days) and combined with 3 µl of Proteinase K, followed by immediate freezing at – 20 °C. After all timepoints were collected, samples were incubated at 65 °C for 30 mins, then at 95 °C for 2 mins, and rapidly cooled to 4 °C over 5 mins. 5 µl of this mixture was then combined with 5 µl 1xPBS and 3 µl loading dye (ThermoFisher #R0611) and run on 8% native PAGE gel.

### Dynamic light scattering

Nanostructures were prepared at 25 µM in 0.5 ml of 1xPBS as previously described and measured for their hydrodynamic diameter on Zetasizer Lab Blue (Malvern Pananalytical) in 0.5-1 ml disposable cuvettes. Measurements were performed using the recommended automatic settings at side-scattering, with at least 5 rounds of measurements, each with 5-10 reads. Two technical replicates were performed per group, and only readouts with Polydispersity index <0.5 were included.

### Animal experiments

All animal procedures were conducted according to the Institutional Animal Care and Use Committee (IACUC) protocols of the University of Massachusetts Chan Medical School (IACUC protocol 202000010) and in accordance with the National Institutes of Health Guide for the Care and Use of Laboratory Animals. FVB/NJ or C57BL/6 female mice (8-14 weeks old) were used for all systemic studies, and male C57BL/6 mice (10-12 weeks old) were used for all brain studies (ICV injections). The colonies were maintained and housed at pathogen-free animal facilities at UMass Chan Medical School with a 12 h light/12 h dark cycle at controlled temperature (23 ± 1 °C) and humidity (50% ± 20%) with free access to food and water.

### siRNA accumulation quantification for plasma clearance and pharmacokinetics

siRNA plasma clearance kinetics were assessed by measuring plasma siRNA concentrations via serial blood microsampling, as previously described.^45,63^ Briefly, mice (*n* = 4) were injected intravenously (20 nmol total siRNA to be quantified), followed by blood microsampling (10–20 μL). The saphenous vein was punctured using a 30G1/2 sterile lancet, and blood drops were collected using Microvette CB300 K2R tubes (Sarstedt, 16.444.100). Blood was collected at the following timepoints: 1 min, 5 min, 15 min, 30 min, 1 h, 2 h, 4 h, 6 h, and 24 h. Blood was then centrifuged for 10 min at 10,000 RPM at 4°C to isolate plasma. Plasma was then lysed (2 μL plasma in 200 μL homogenizing buffer [Affymetrix] containing 0.2 mg mL^−1^ proteinase K [Invitrogen]) at 65°C for 30 min. siRNA quantification in plasma was performed using a PNA-hybridization assay as previously described.^63^ The probe used to hybridize to the HTT antisense strand of the HTT siRNA was purchased from PNAbio and had the following sequence: Alexa 488-OO-AAATATCAGTAAAGAGATTAA. DCA-siRNA plasma circulation data was conducted in the same study, but was previously reported in a separate publication by Fakih et. al.^45^ For the study comparing FMRN to DCA at one-month post injection, we used a PNA probe that hybridize to the JAK1 antisense strand of the JAK1 siRNA carried by all groups, with the following sequence: Alexa488-OO-GTACAAAGGAATCTGA-KK as previously reported.^13^

### *In vivo* efficacy following systemic injection

siRNAs and structures formulated in 1x PBS were prepared at a 200 µM concentration as described above. The siRNAs or structures were then administered subcutaneously between the shoulder blades of mice or intravenously via the tail vein. The dosages indicated for each experiment (e.g., 20 mg/kg) were based on siRNA dose. For example, for the T-3xTarget group, the 20 mg/kg dose is 20 mg/kg per siRNA target (*Htt*, *Jak1*, and *Msh3*). For T-3xHTT, it is also a total of 20 mg/kg HTT-siRNA dose, but it is a 6.67 mg/kg dose of the structure as it carries three copies of the same HTT siRNA. Animals were then euthanized at the indicated timepoints, and tissues were harvested and processed for mRNA quantification and siRNA accumulation as described above.

### *In vivo* efficacy following stereotactic intracerebroventricular (ICV) injections

ICV injections were performed as previously described.^28,64^ all siRNA groups were first prepared in water, then they were dried and resuspended in artificial CSF (aCSF, 137 mM NaCl, 5 mM KCl, 20 mM glucose, 8 mM HEPES, 2.3 mM CaCl_2_, 1.3 mM MgCl_2_, pH 7.4, supplemented up to 14 mM CaCl_2_, 2 mM MgCl_2_ ^65^). Briefly, mice (Jackson Laboratory, Strain # 005304) aged 10-12 weeks were anesthetized with 2.5% isoflurane in oxygen. Hair at surgical sites was shaved with an electric razor, and a sterile surgical field was set up around the animal’s head. A burr was used to drill a small hole at the following coordinates relative to bregma: −0.2 mm posterior, 1.0 mm mediolaterally. The needle was then placed at a depth of −2.5 mm ventrally. A total volume of 5 μL was injected per ventricle at a rate of 750 nl/min. Mice were monitored post-injection until sternal. All doses are based on molar amounts of active siRNA as described above.

At endpoint, the brain was harvested and placed in a brain matrix (Braintree Scientific). A 1-mm slice was then taken from the regions of interest, and 1.5 × 1.5-mm punches were taken and placed in RNALater (935 ml MilliQ water, 700 g ammonium sulfate, 25 ml 1M Sodium Citrate, 40 ml 0.5 M EDTA, pH 5.2 using H_2_SO_4_) for 24 h at 4°C, then frozen at −80°C until processing for subsequent experiments. Biopsies were dried gently, weighed, and added to 500 μL of homogenizing solution (Invitrogen; QS0517) containing 0.2 mg/mL proteinase K (Invitrogen, #AM3546), in a QIAGEN collection microtube holding a 3-mm tungsten bead. The tissues were then homogenized for 10 min at 25 Hz using a QIAGEN TissueLyser II, followed by incubation at 55 °C for 30 min. Lysate was then used as described above for mRNA quantification, or for siRNA accumulation (described below), or stored at -20 °C.

### Lipoprotein SEC

For lipoprotein profiling, we followed the same protocol previously described.^44,60^ Briefly, mice were injected s.c. with 20 nmol siRNA dose (equivalent to10 mg/kg siRNA dose) labeled with Cy3.For DCA, the sense strand is labeled with Cy3, whereas for the FMRNs, one of the Arms is labeled with Cy3 (Htt-carying Arm strand, as shown in the supplementary table 1). After 30 mins, whole mouse blood (∼500 μL) was collected in a sterile ethylenediaminetetraacetic acid (EDTA)-coated tube following cheek incision with a lancet. Samples were spun at 10,000 rpm for 10 min at 4°C. Fifty microliters of plasma was directly injected on a Superose 6 Increase 10/300 size exclusion column (GE Healthcare). Oligonucleotide migration was monitored at 570 nm, and lipoprotein protein content was monitored by absorbance at 280 nm.

### siRNA tissue accumulation quantification

To quantify siRNA accumulated in tissues following systemic administration, PNA hybridization assay was used on the lysate prepared for mRNA quantification, as previously described, with slight modification.^63,64^ Samples were weighed to quantify siRNA accumulation per milligram of tissue. The accumulation was measured using a custom Alexa488-labeled fluorescent PNA oligonucleotide probe (Alexa 488-OO-AAATATCAGTAAAGAGATTAA, PNABio) that is fully complementary to the antisense strand of the HTT-targeting siRNA. The prepared lysate of each tissue was first treated with 50 μL of 3 M potassium chloride to precipitate any sodium dodecyl sulphate present, followed by centrifugation for 15 min at 5000×g. siRNAs in the clear supernatant were then collected and hybridized to the PNA probe at 95 °C, followed by slow cool down. Anion-exchanged chromatography was used to analyze the sample mixtures on an Agilent 1260 Infinity quad-pump HPLC with a 1260 FLD fluorescent detector, and quantification was based on comparison to standard curves of spike-in siRNA in tissue lysate as previously described.

### Quantification of myostatin protein in plasma

Blood samples were collected from control or siRNA injected animals via cheek bleeding, where 20–50 µl was collected on the indicated times (weekly). Blood was collected in ethylenediaminetetraacetic acid (EDTA) coated tubes, and then spun at 10 000 rpm for 10 mins at 4°C to separate plasma. Plasma was collected and stored at −80°C until ready to use for quantification. *MSTN* protein was quantified using ELISA kits (Bio-Techne/R&D systems, #DGDF80) as previously described^48^ where 3 µl of plasma was used per sample and following standard provided protocol.

### Blood diagnostics

Blood sampling for blood diagnostics was performed at 72h by collecting blood via cheek bleed. In total, 200 μL of blood was collected in a lithium heparin-coated BD Microtainer tube (BD, #365965) for blood chemistry test. 100 μL of blood was collected in a K2 EDTA-coated bd Microtainer tube (BD, #365974) for a complete blood count (CBC) test. Blood chemistry and CBC diagnostics were conducted by the Diagnostics Laboratory in the Department of Animal Medicine at UMass Chan Medical School.

### Graphs and statistical analyses

Data were analyzed using GraphPad Prism 10.1.2 software for Windows (GraphPad Software, Inc., San Diego, CA). For each independent experiment, the levels of human or mouse target mRNA silencing were normalized to the mean of the untreated group (for *in vitro* screening) or the NTC group (for *in vivo* studies). *In vivo* data were analyzed using a one-way ANOVA with a post hoc Dunnett multiple comparison test when comparing to the control group only, or Tukey’s multiple comparison test when comparing all groups to each other. Asterisks (*) denote gene expression significance relative to NTC or indicated group with bracket (\**p* <0.05, \*\**p* <0.01, \*\*\**p* <0.001, \*\*\*\**p* <0.0001), non-significant was not indicated. Graphs are plotted as mean ± standard deviation. Further statistical details are provided in each figure caption for each study.

## CONTRIBUTIONS

HHF and AK conceived the project. HHF, QT, AS and AK contributed to the experimental design. HHF, AS, QT, RG, BMB, RCF, KYG and DE contributed experimentally. HHF and HS contributed to the initial structural design of nanostructures. HHF, RG, BMB and DE contributed to compound synthesis. HHF and AK wrote the manuscript. All authors provided feedback and approved of the manuscript.

## CORRESPONDING AUTHOR

<u>Hassan H. Fakih and Anastasia Khvorova</u>.

## DATA AVAILABILITY

All data presented in the main text and the supplementary information are available from the corresponding authors upon request.

## ACKNOWLEDGEMENTS

We thank Khvorova lab members (past and present) for insightful discussions and support, particularly Ken Yamada and Samuel Hildebrand. We thank the animal medicine core at UMass Chan Medical School, for their help and support on multiple studies. We also thank Emily Haberlin for her assistance with writing the manuscript. All figures were created with PowerPoint and BioRender.com.

## SUPPLEMENTARY INFORMATION

Supplementary Fig. S1– 7 and Supplementary Tables S1.

---

**Extended Figure 1:**
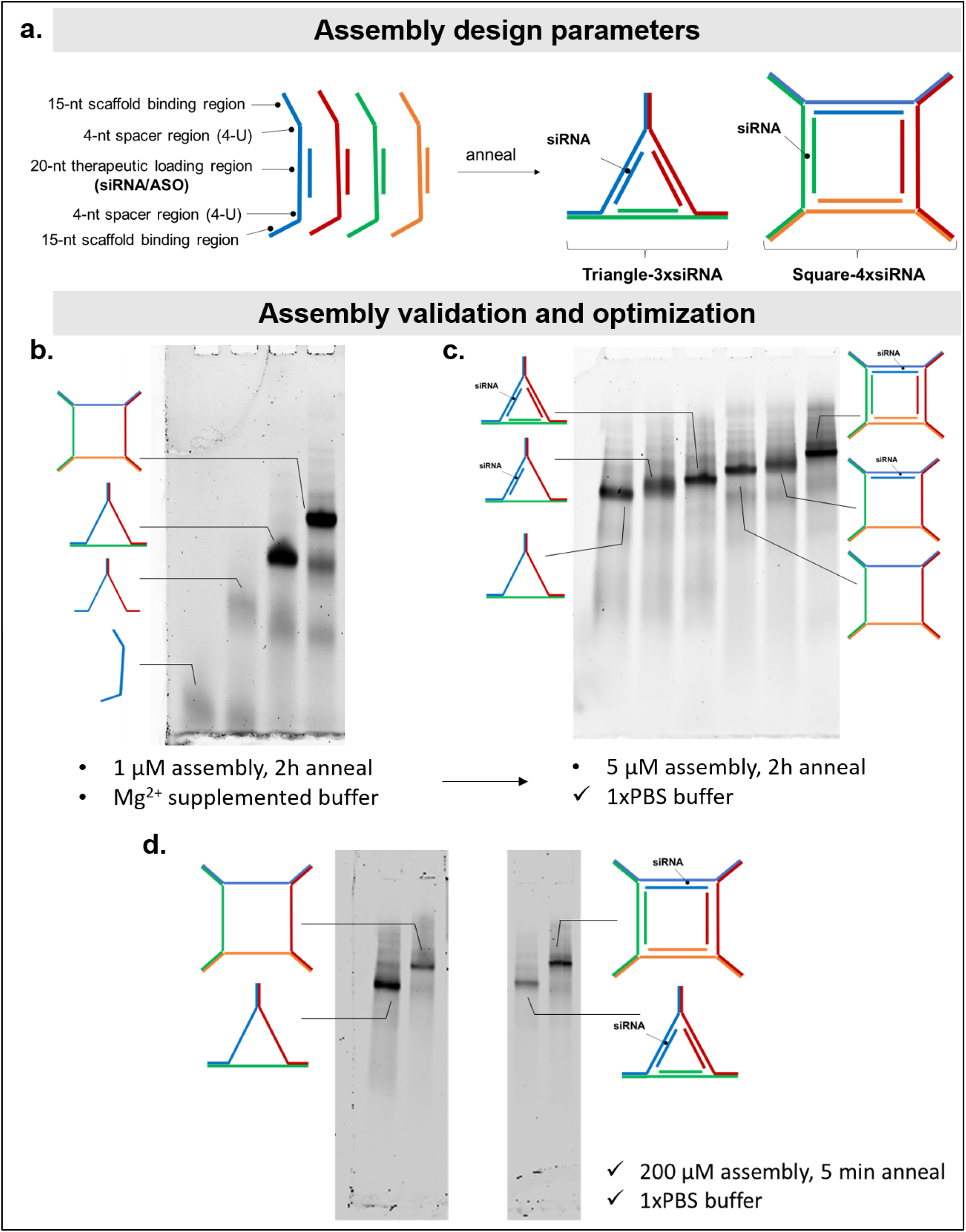
Design, assembly, and structural characterization of FMRNs architectures. **a.** Schematic representation of FMRN architectural parameters. Each polygon arm is composed of three functional domains: (i) 15-nt (or 20-nt) scaffolding regions at the 5’ and 3’ termini that facilitate inter-strand hybridization via “sticky ends”; (ii) a central 20-nt therapeutic region designed to incorporate the antisense strand of an siRNA or ASO; and (iii) 4-nt single-stranded flexible vertices flanking the therapeutic domain to enable precise folding. Self-assembly of three or four arm strands (different colors for different sequences) yields triangular (T-FMRN) or square (S-FMRN) polygons with stochiometric loading of 3 or 4 siRNAs, respectively. **b.** Step-wise electrophoretic characterization of T- and S-FMRN assembly in magnesium-supplemented TBE buffer. Sequential addition of arm strands results in distinct bands with progressively lower electrophoretic mobility, confirming monodisperse assembly (lanes 3 and 4). Assembly was conducted using a standard 2-h slow-annealing cycle. **c.** Assembly in phosphate-buffered saline (PBS) and characterization of payload loading. Electrophoretic characterization demonstrates a gradual decrease in mobility corresponding to the stochiometric loading of siRNA copies (1 vs. 3 or 4). **d.** Scalability and kinetic optimization of FMRN assembly. Gel confirms robust, monodisperse assembly of T- and S-FMRNs (with or without siRNA payloads) at *in vivo*-relevant concentrations (100 µM siRNA equivalent, 33.34 µM T-3xsiRNA equivalent, and 25 µM S-4xsiRNA equivalent) in PBS. Rapid assembly (5-min “quick-anneal” at 95 °C followed by rapid cooling to 4 °C) yields structural integrity comparable to the 2h anneal. (All assemblies were analyzed at 1 µM unless otherwise specified. Electrophoresis was performed on 8% non-denaturing PAGE gels at 130 V for 1 h in 1× TBE buffer).

**Supplementary Figure 1:**
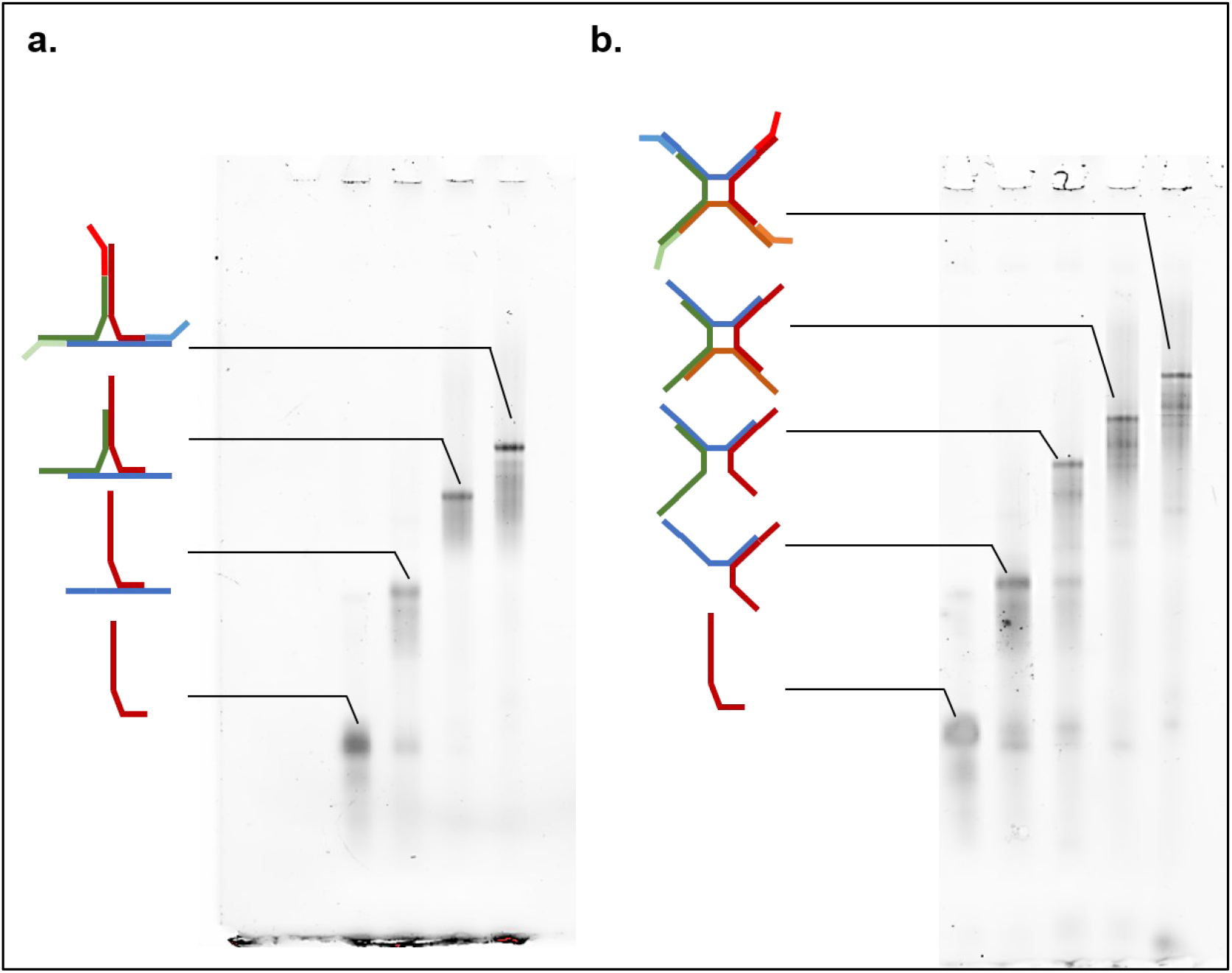
Stepwise assembly of T- and S-FMRN variant 4. **a.** T-FMRN and **b.** S-FMRN variant 4 assembly validation in step-wise assembly gel, adding one arm at a time, and then all antisense strands to complete the siRNA loading. Assembly was at 10 µM in 1xPBS, annealed for 5 mins at 95C followed by cooling at room temperature. Electrophoresis was performed on 8% non-denaturing PAGE gels at 130 V for 1 h in 1× TBE buffer.

**Supplementary Figure 2:**
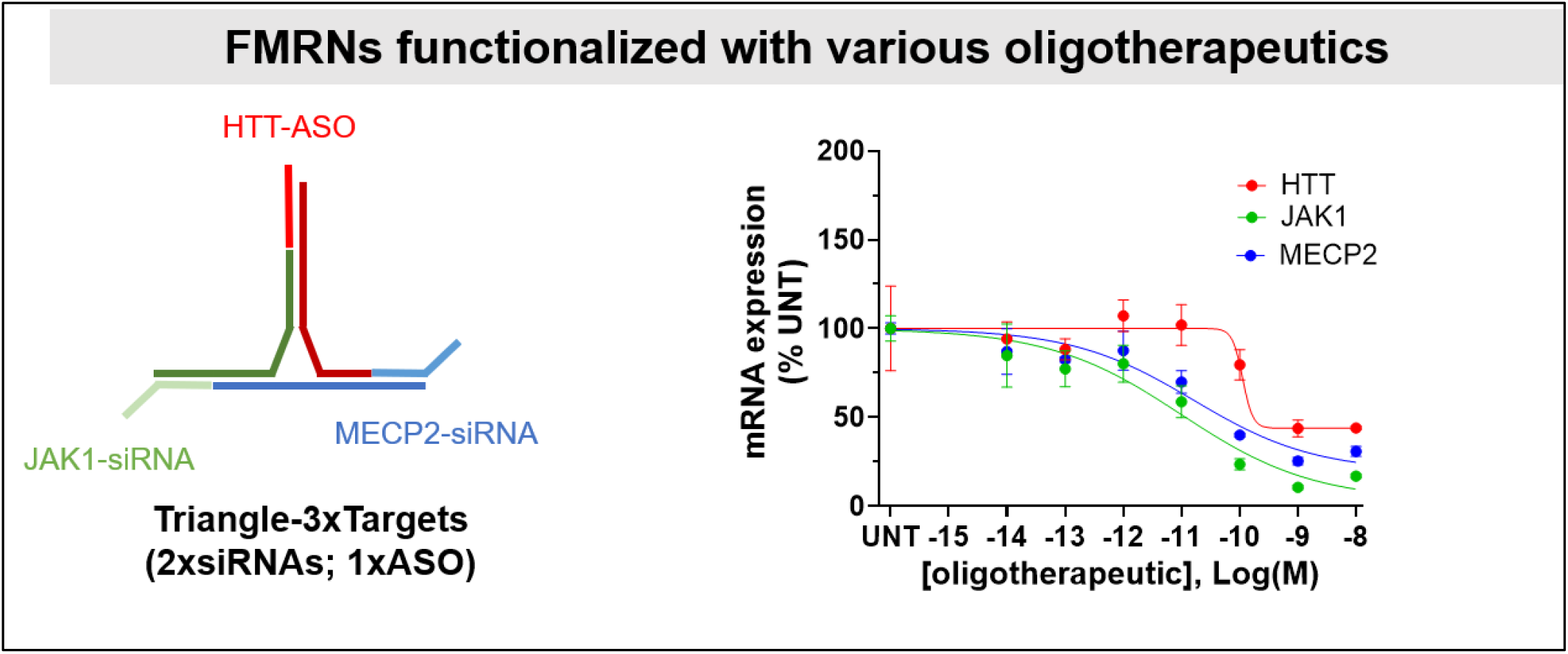
Multi-modality functionalization of T-FMRN, by delivering one ASO targeting *HTT* and two siRNAs targeting *JAK1* and *MECP2* mRNAs. Dose-dependent siRNA efficacy *in vitro* against each target is observed. (study was conducted in HeLa cells, with siRNAs transfected at the indicated concentrations for 72h ; n=3/group; mRNA quantification via Quantigene 2.0 assay)

**Supplementary Figure 3:**
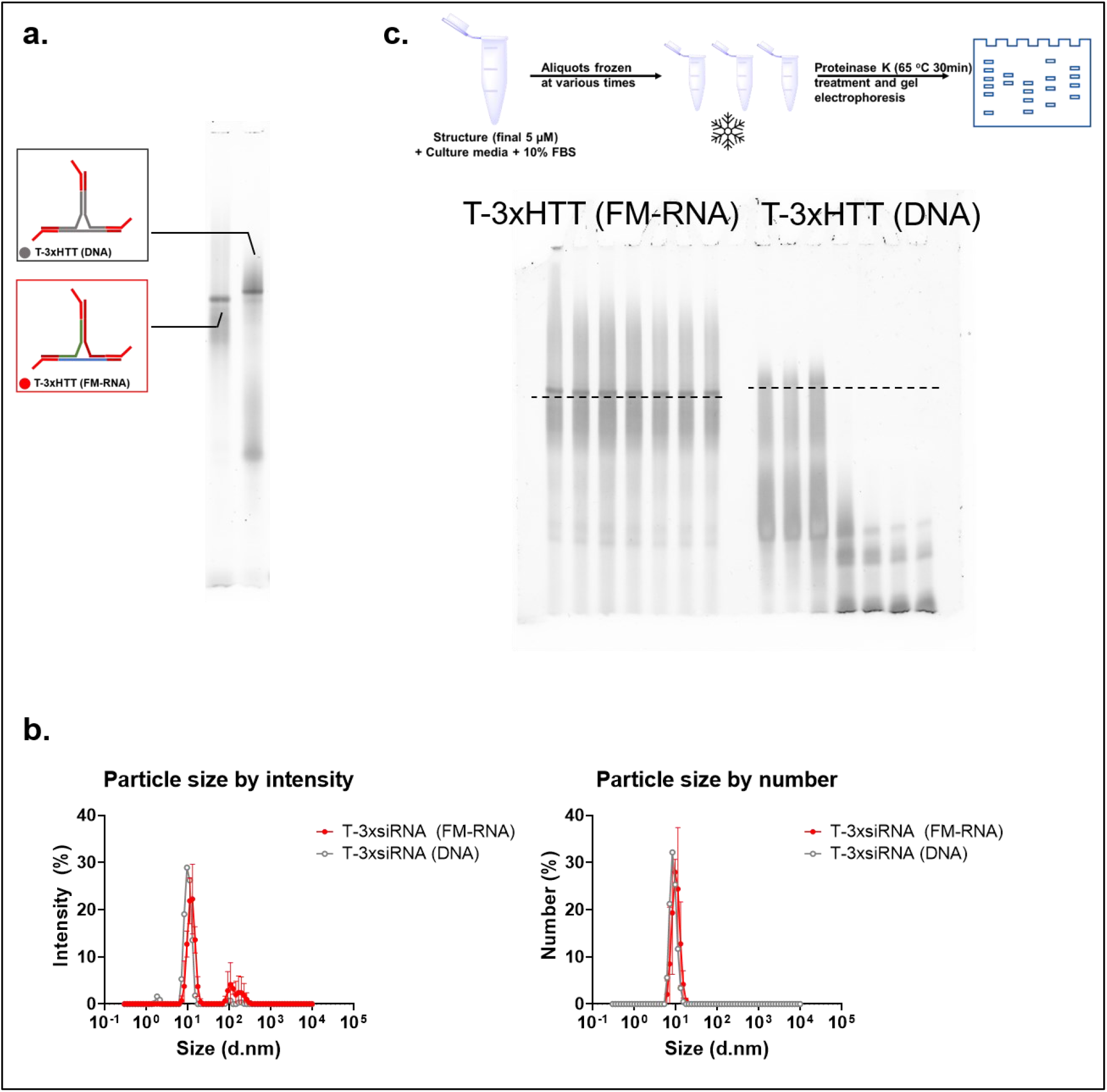
Physicochemical characterization of Triangle-3xsiRNA as fully modified RNA or DNA-based scaffold. **a.** Native PAGE (8%) analysis showing assembly of both T-3xHTT (FMRN) and T-3xHTT (DNA) constructs showing complete formation, with FMRN structure forming slightly better likely due to high melting temperature and stronger binding. **b.** Hydrodynamic size distribution measured by dynamic light scattering (DLS), represented as intensity and number percentages. **c.** Native PAGE (8%) gel of the nuclease stability study as detailed in the schematic. (Electrophoresis was performed on 8% non-denaturing PAGE gels at 130 V for 1 h in 1× TBE buffer. DLS measurements are n=2 technical replicates, with concentrations of FMRNs at 25uM; data represented as mean ± S.D.; graphical elements created with Biorender; Fakih et. al.).

**Supplementary Figure 4:**
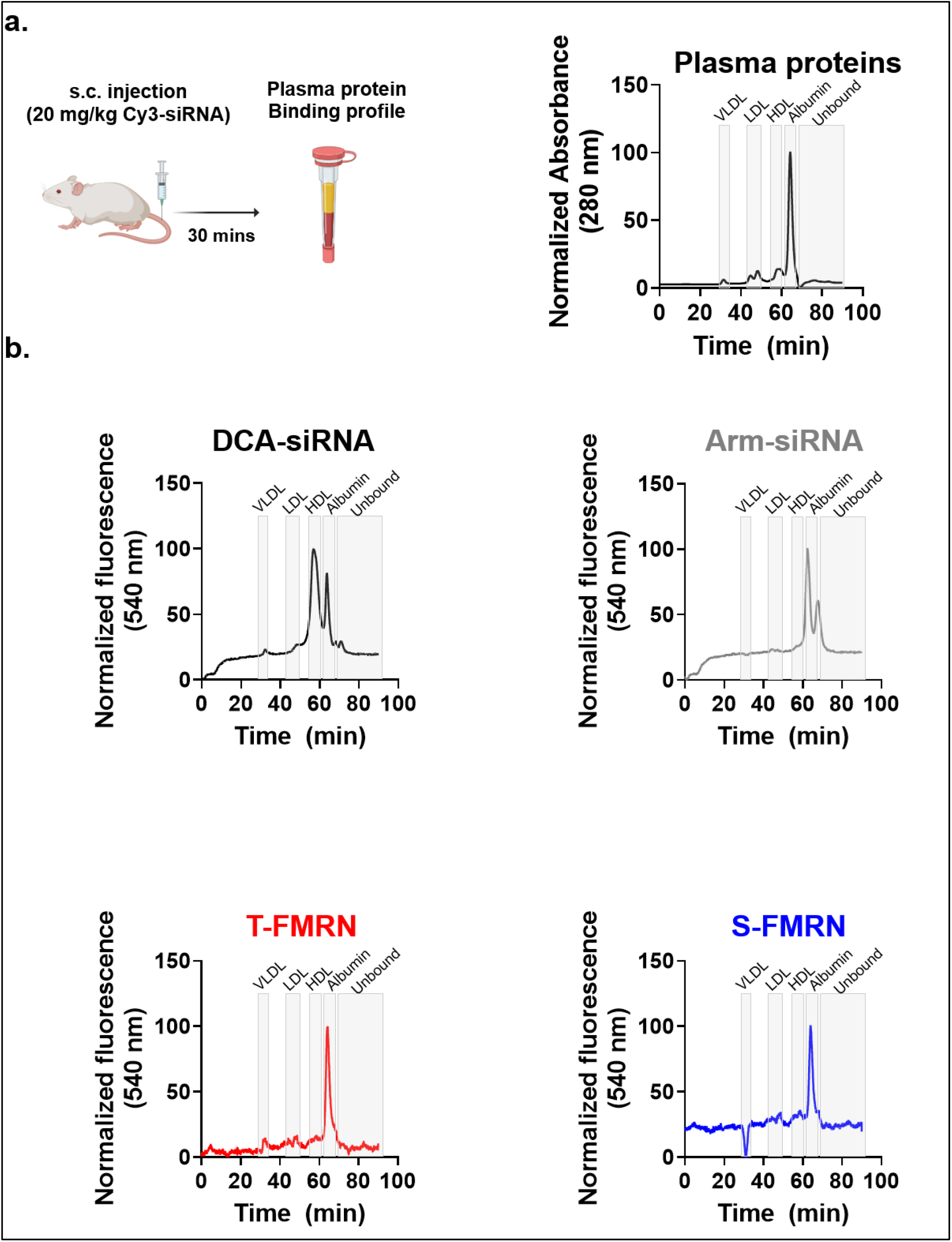
Characterization of the plasma protein binding profile of FMRN reveals albumin-only binding. **a.** Study schematic, where Cy3-labeled constructs were injected subcutaneously at 20 mg/kg siRNA dose, and plasma collected 30 mins post injection followed by analysis via size-exclusion chromatography (SEC) as previously reported (Fakih et. al. MTNA 2023). **b.** Plasma protein binding profile of each construct (DCA-, Arm-, T-, and S-siRNAs), proteins monitored and assigned at 280 nm absorbance and constructs binding profile monitored via fluorescence of Cy3 at 570 nm. Elution signals were normalized to the highest peak. (n=2 mice/group.; graphical elements created with Biorender; Fakih et. al.).

**Supplementary Figure 5:**
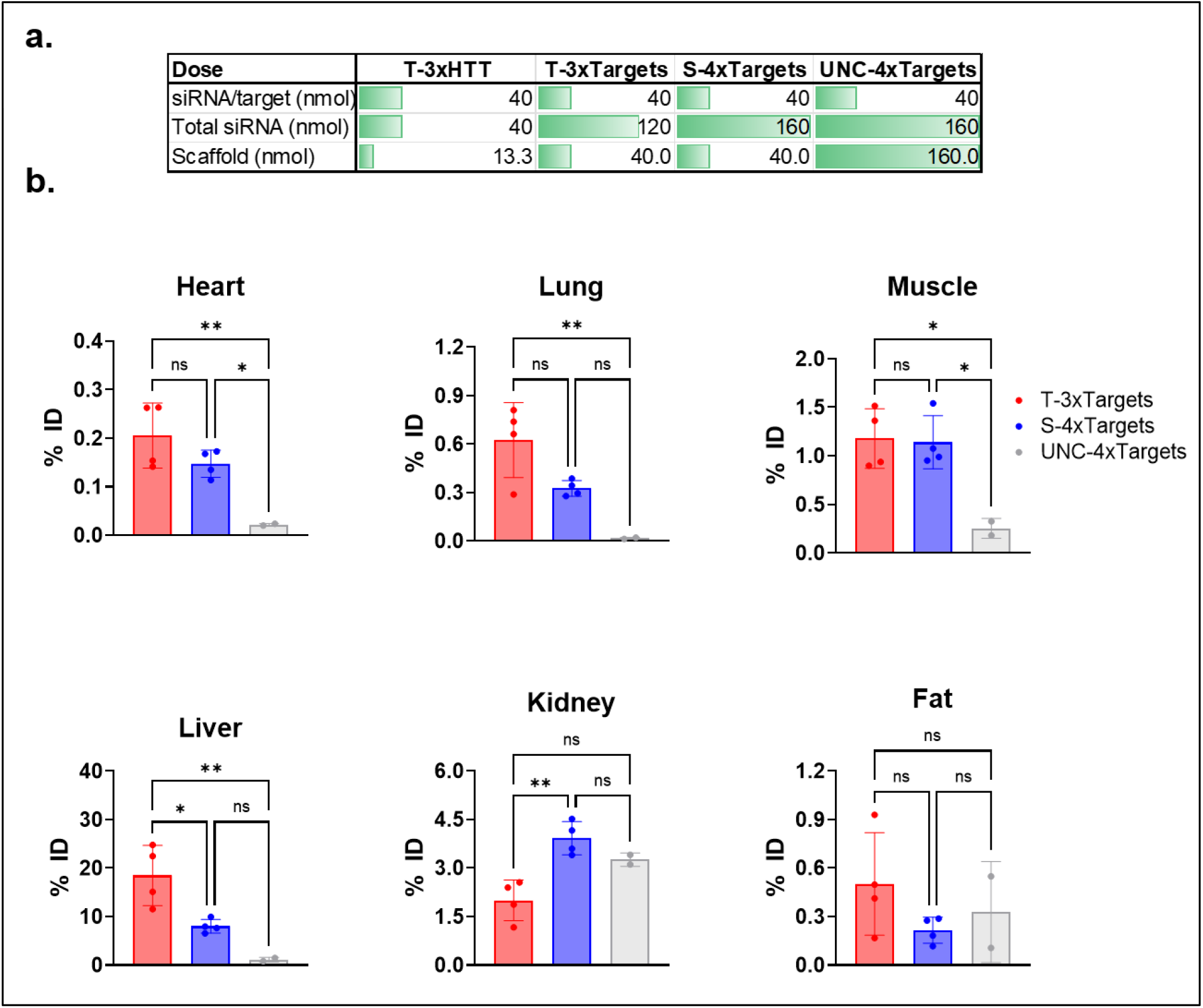
Dose information for the FMRNs and percentage of injected dose (%ID) accumulated in each tissue. **a.** Dose calculation in nmol for each of the FMRNs used across figure 2 and Figure 4, given in nmol of each siRNA/target, total nmol of siRNA, and nmol of scaffold. 40 nmol siRNA is equivalent to 20 mg/kg in these studies. **b.** %ID for each group in each tissue or organ measured, based on the quantification of siRNA accumulation in the corresponding figures (measuring the HTT-siRNA accumulation and %ID based on the 40nmol dose of that siRNA); graphical elements created with Biorender; Fakih et. al.).

**Supplementary Figure 6:**
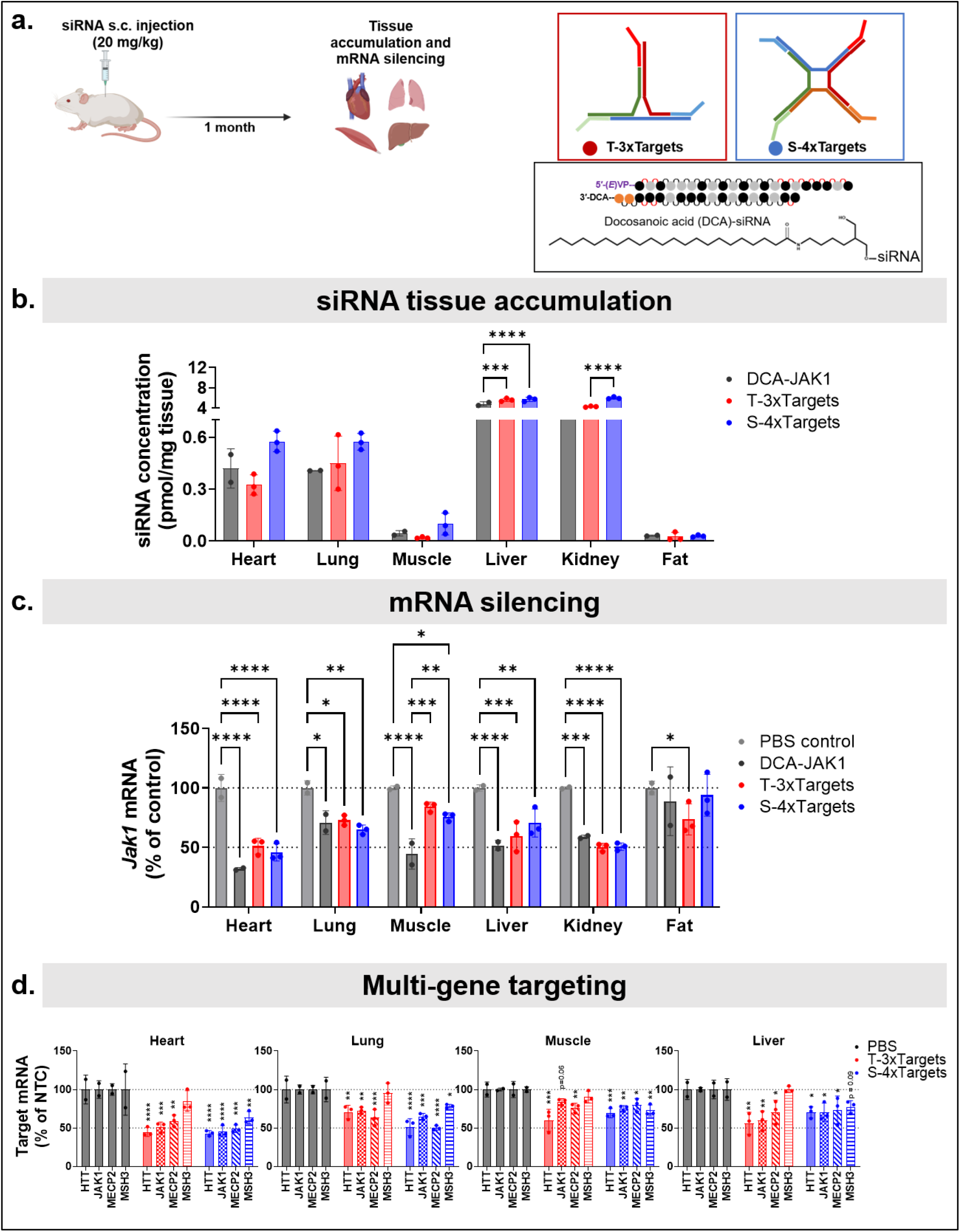
T- and S-FMRN achieve durable siRNA tissue accumulation and target mRNA silencing that is comparable to lipid conjugated DCA-siRNA at one-month post injection, with multi-gene targeting in various extra-hepatic tissues. **a.** Scheme of *in vivo* study evaluating the distribution and efficacy of FMRN at one-month post injection, in comparison to DCA-siRNA. Mice received subcutaneous (s.c.) injections of either PBS control, DCA-JAK1, T-3xTargets or S-4xTargets, (40 nmol siRNA dose/target, equivalent to 40 nmol scaffold dose for all groups). Tissues were harvested one month post injection to assess siRNA accumulation and mRNA silencing of JAK1 for all groups, and HTT, JAK1, MSH3, and MECP2 mRNAs for FMRNs groups. **b.** siRNA Tissue accumulation and **c.** JAK1 mRNA silencing profiles across major extrahepatic tissues. **d** Simultaneous multi-gene target silencing at one month. mRNA expression levels for HTT, JAK1, MECP2, and MSH3 were compared between the T-3xTargets and S-4xTargets architecture in heart, lung, muscle and liver. All study groups are *n* = 2-3 mice/group. Statistical analysis was performed in GraphPad Prism: for c. one-way ANOVA analysis followed by Tukey’s multiple comparisons across all groups for HTT silencing; for d. two-way ANOVA analysis followed by Dunnet’s multiple comparisons comparing PBS control for every target mRNA expression; non significant comparisons not shown, \**p* < .05, \*\**p* < .01, \*\*\**p* < .001, \*\*\*\**p* < .0001. (tissue accumulation measured by PNA assay; mRNA measured by the Quantigene 2.0 Assay; data represented as mean ± S.D.; dotted line on all graphs represents 100% and 50% remaining mRNA expression; graphical elements created with Biorender; Fakih et. al.).

**Supplementary Figure 7:**
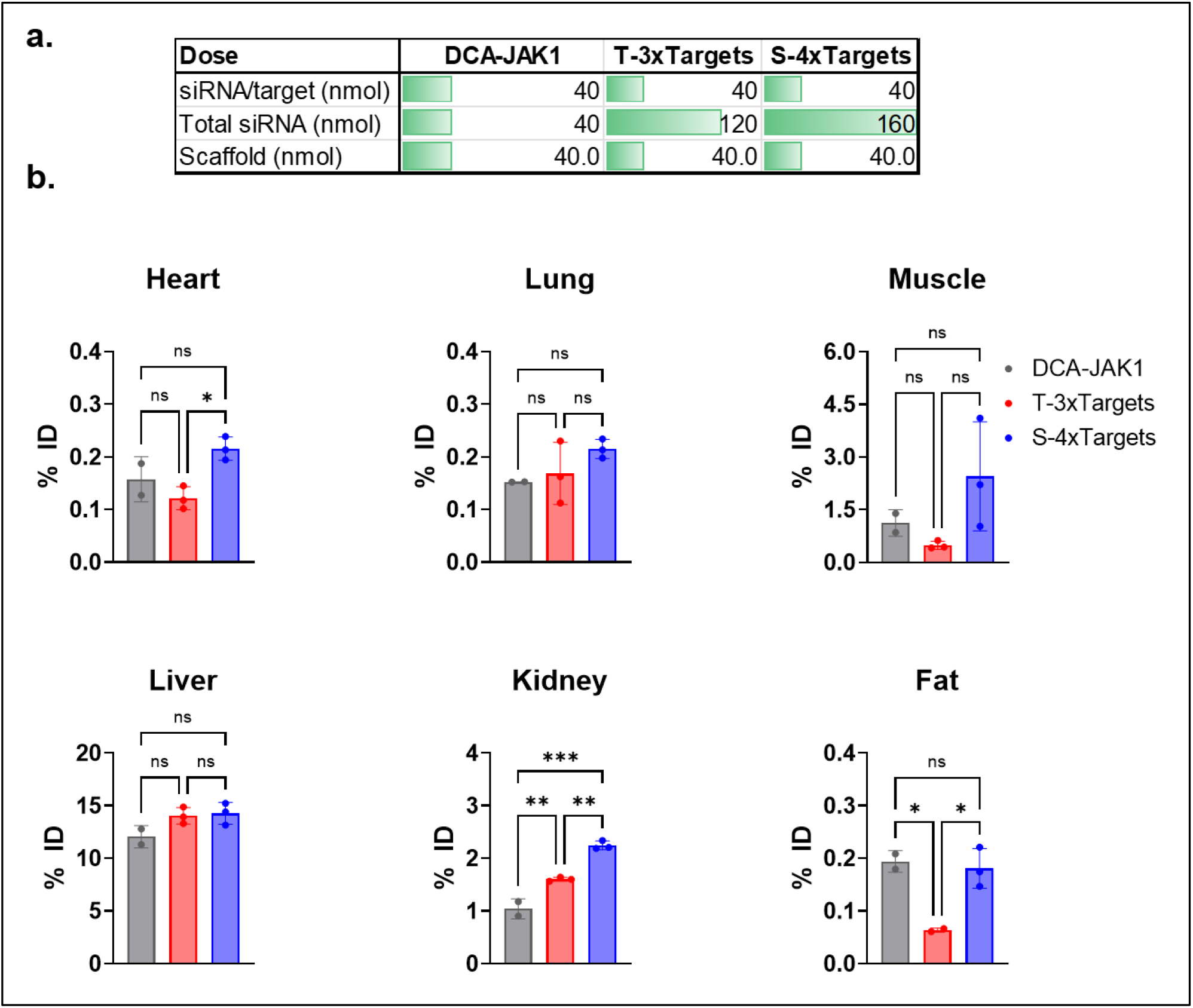
Dose information for the FMRNs and percentage of injected dose (%ID) accumulated in each tissue. **a.** Dose calculation in nmol for each of the FMRNs from the study comparing the durability of the FMRNs to that of DCA-siRNA (Supplementary figure 6), given in nmol of each siRNA/target, total nmol of siRNA, and nmol of scaffold. 40 nmol siRNA is equivalent to 20 mg/kg in these studies. **b.** %ID for each group in each tissue or organ measured, based on the quantification of siRNA accumulation in the corresponding figures (measuring the JAK1-siRNA accumulation and %ID based on the 40nmol dose of that siRNA); graphical elements created with Biorender; Fakih et. al.).

**Supplementary Figure 8: T.**
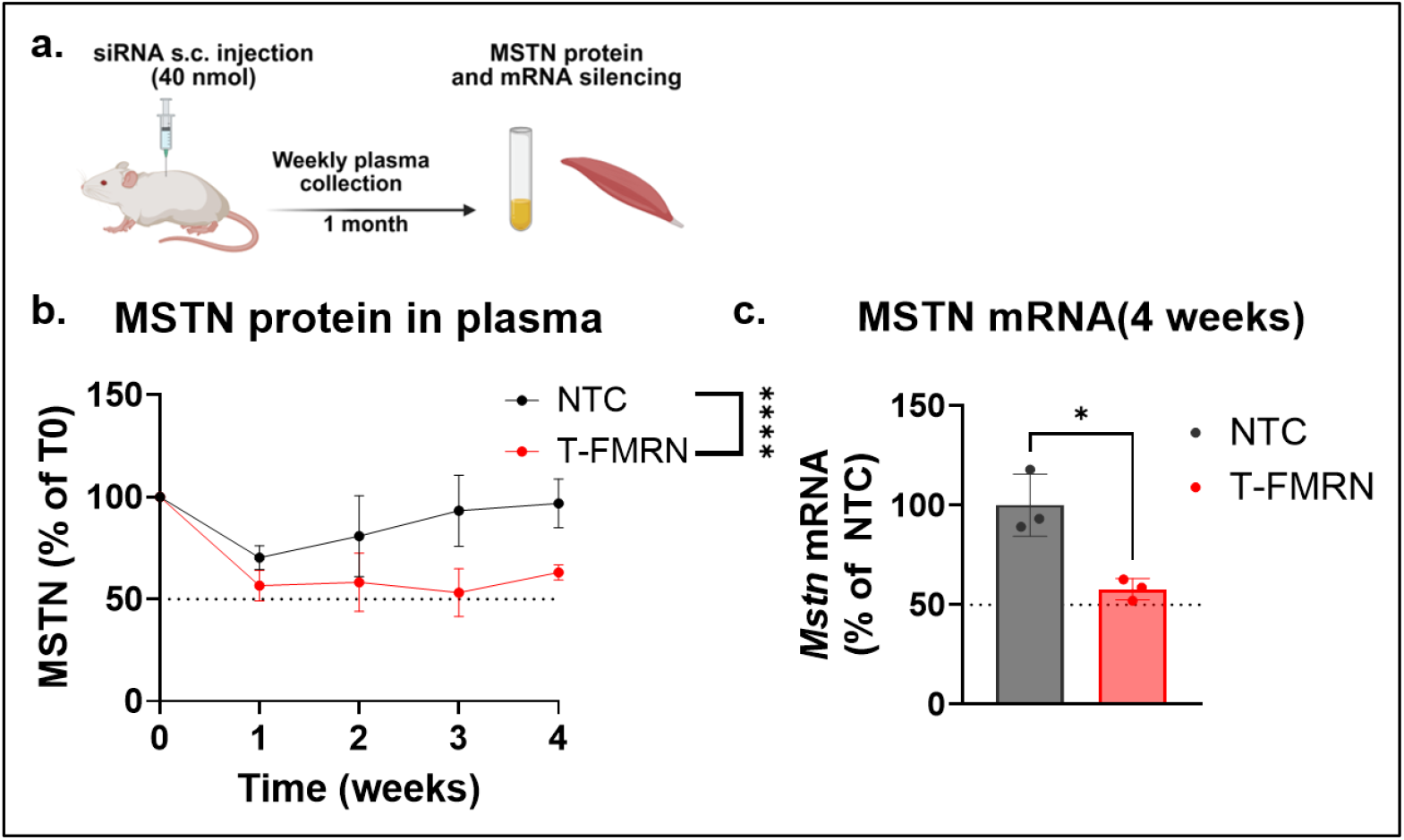
-FMRN supports potent and durable silencing of myostatin in muscle at the mRNA and protein level. **a.** Scheme of *in vivo* study evaluating the longtuidnial silencing of MSTN in muscle by T-FMRN at the mRNA and protein level. Mice (n=3/group) received subcutaneous (s.c.) injections of either NTC control or T-FMRN loaded with siRNA targeting Mstn. (40nmol siRNA dose, equivalent to 40 nmol scaffold dose). Plasma was collected weekly to measure MSTN levels, and muscle tissue collect at the end of study on fourth week. **b.** MSTN protein and **c.** Mstn mRNA levels. Measured at the indicated timepoints Statistical analysis was performed in GraphPad Prism: two-way ANOVA analysis for MSTN protein data; T-test analysis for Mstn mRNA expression; non significant comparisons not shown, \**p* < .05, \*\**p* < .01, \*\*\**p* < .001, \*\*\*\**p* < .0001. (Protein levels measured by ELISA assay; mRNA measured by the Quantigene 2.0 Assay; data represented as mean ± S.D.; dotted line on all graphs represents 50% remaining protein or mRNA expression; graphical elements created with Biorender; Fakih et. al.).

**Supplementary Figure 9.**
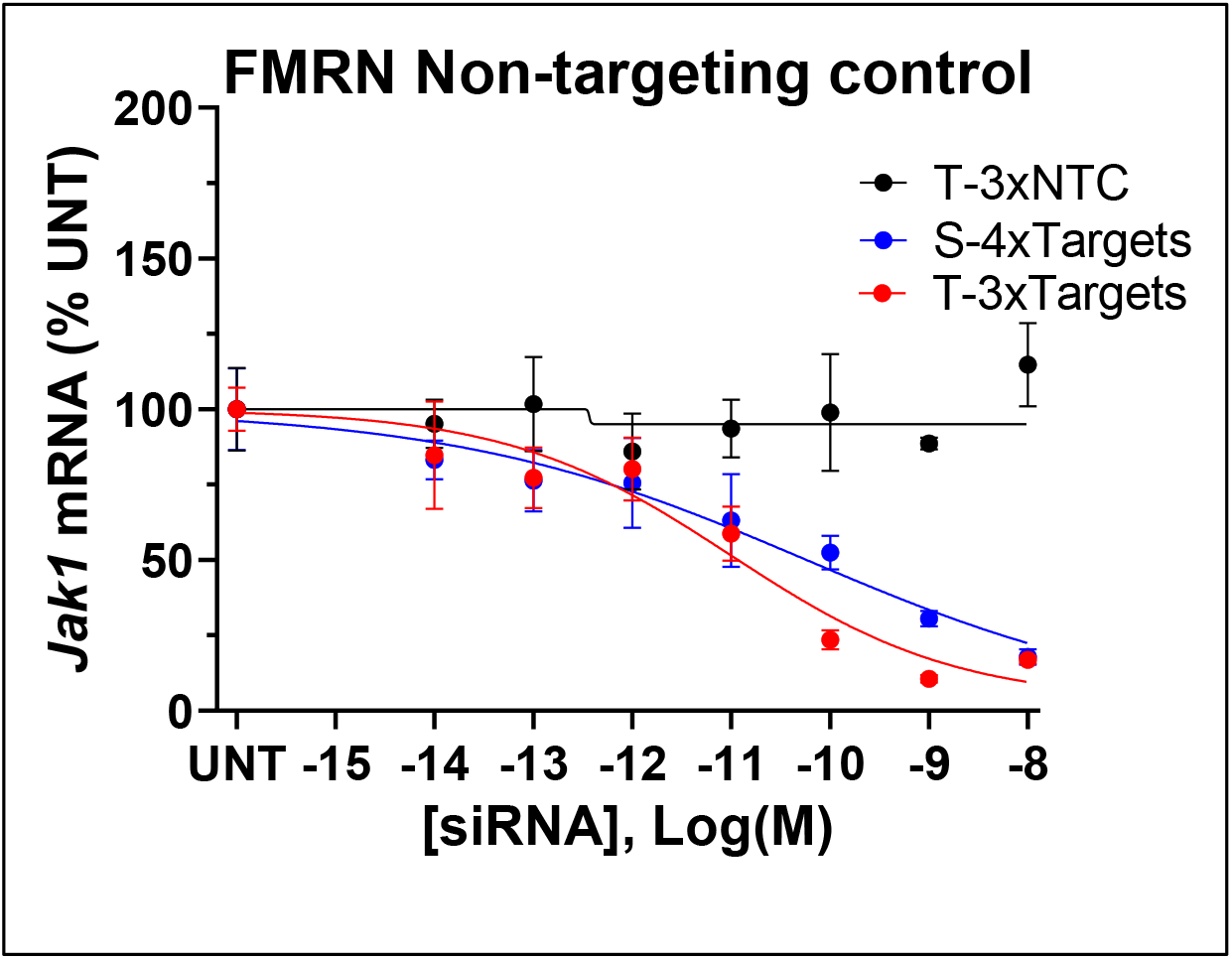
T-FMRN loaded with 3 siRNA NTCs does not induce gene silencing *in vitro*. Dose-dependent siRNA efficacy *in vitro* against JAk1 mRNA, comparing T-3xNTC to T-3xTargets S-4xTargets that silence JAK1, to assess if FMRN structural scaffold has any off-target effects. (study was conducted in HeLa cells, with siRNAs transfected at the indicated concentrations for 72h ; n=3/group; mRNA quantification via Quantigene 2.0 assay. The T-3xTargets JAK1 silencing data is identical to that in supplementary figure 2)

**Supplementary Figure 10:**
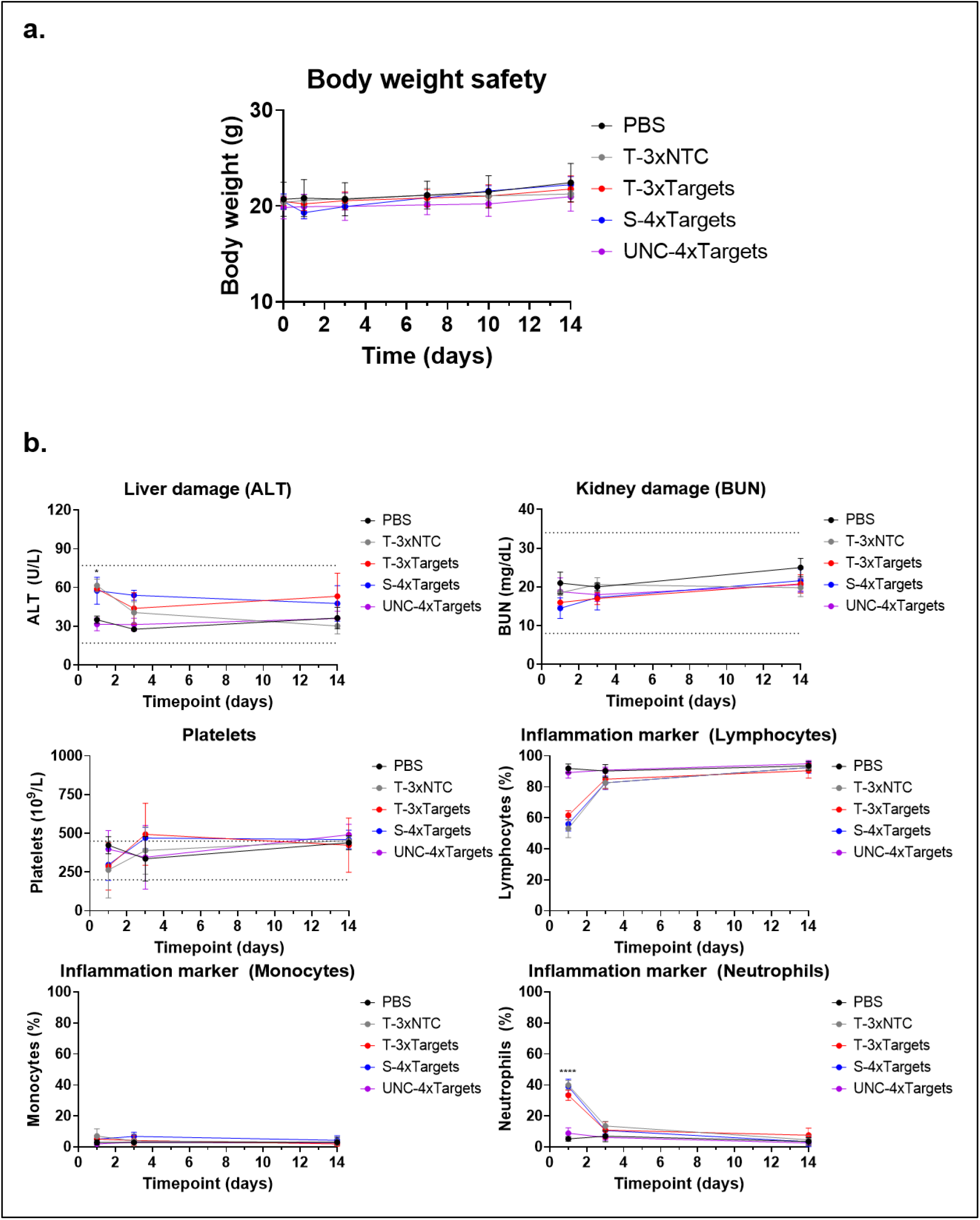
Acute and chronic safety profile of FM-RN following systemic injection. **A.** Body weight of mice tracked over time (preinjection, day 1, 3, 7, 10,14) Markers of liver (ALT) and kidney (BUN) damage, as well as platelet count and immune cell population show no abnormal changes beyond the normal ranges (n=2-5 mice/group.; blood analyzed day 1, 3, and 14 post subcutaneous injection of 40 nmol siRNA/target dose; 40 nmol/scaffold for FMRNs and 160 nmol/scaffold for UNC-siRNA)

**Supplementary Figure 11:**
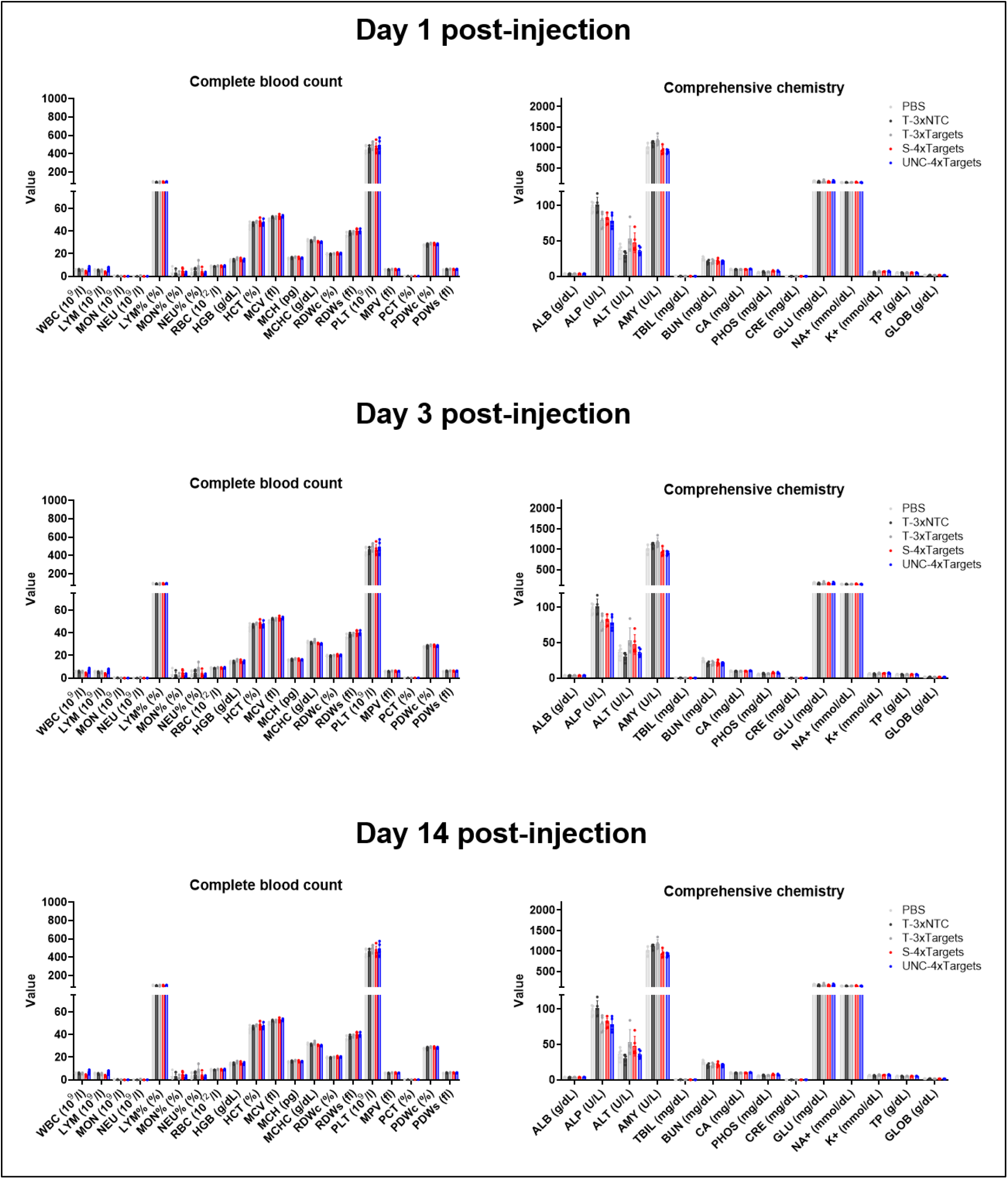
Complete data set of the acute and chronic safety profile of FM-RN following systemic injection.. Complete data set from CBC and blood chemistry measurements. (n=2-5 mice/group.; blood analyzed day 1, 3, and 14 post subcutaneous injection of 40 nmol siRNA/target dose; 40 nmol/scaffold for FMRNs and 160 nmol/scaffold for UNC-siRNA)

**Supplementary Figure 12:**
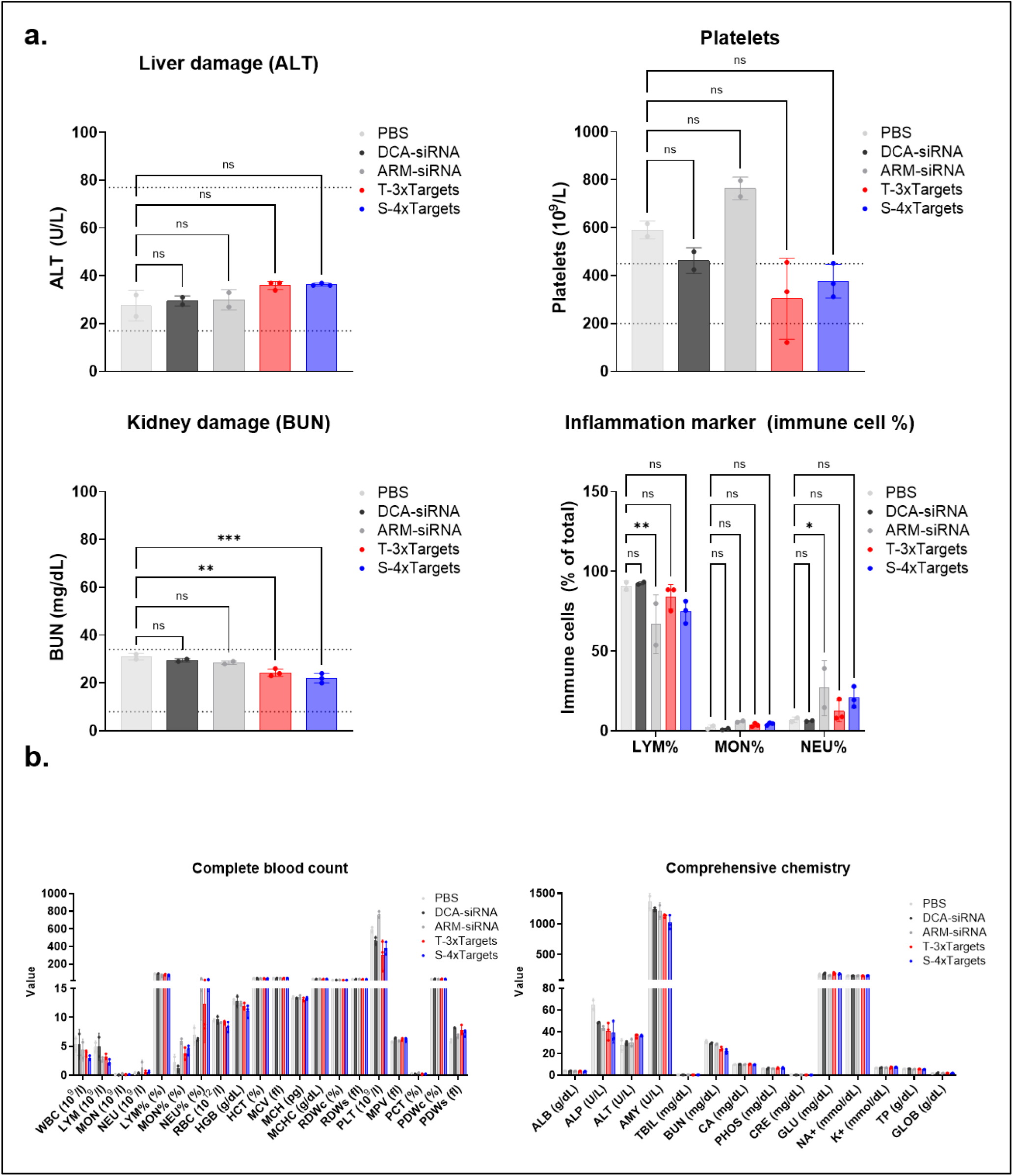
Safety profile of FMRN following systemic injection. **A.** Markers of liver (ALT) and kidney (BUN) damage, as well as platelet count and immune cell population show no abnormal changes. **b.** Complete data set from CBC and blood chemistry measurements. (n=2-3 mice/group.; blood analyzed 72h post subcutaneous injection of 20 mg/kg total siRNA dose)

**Supplementary Figure 13:**
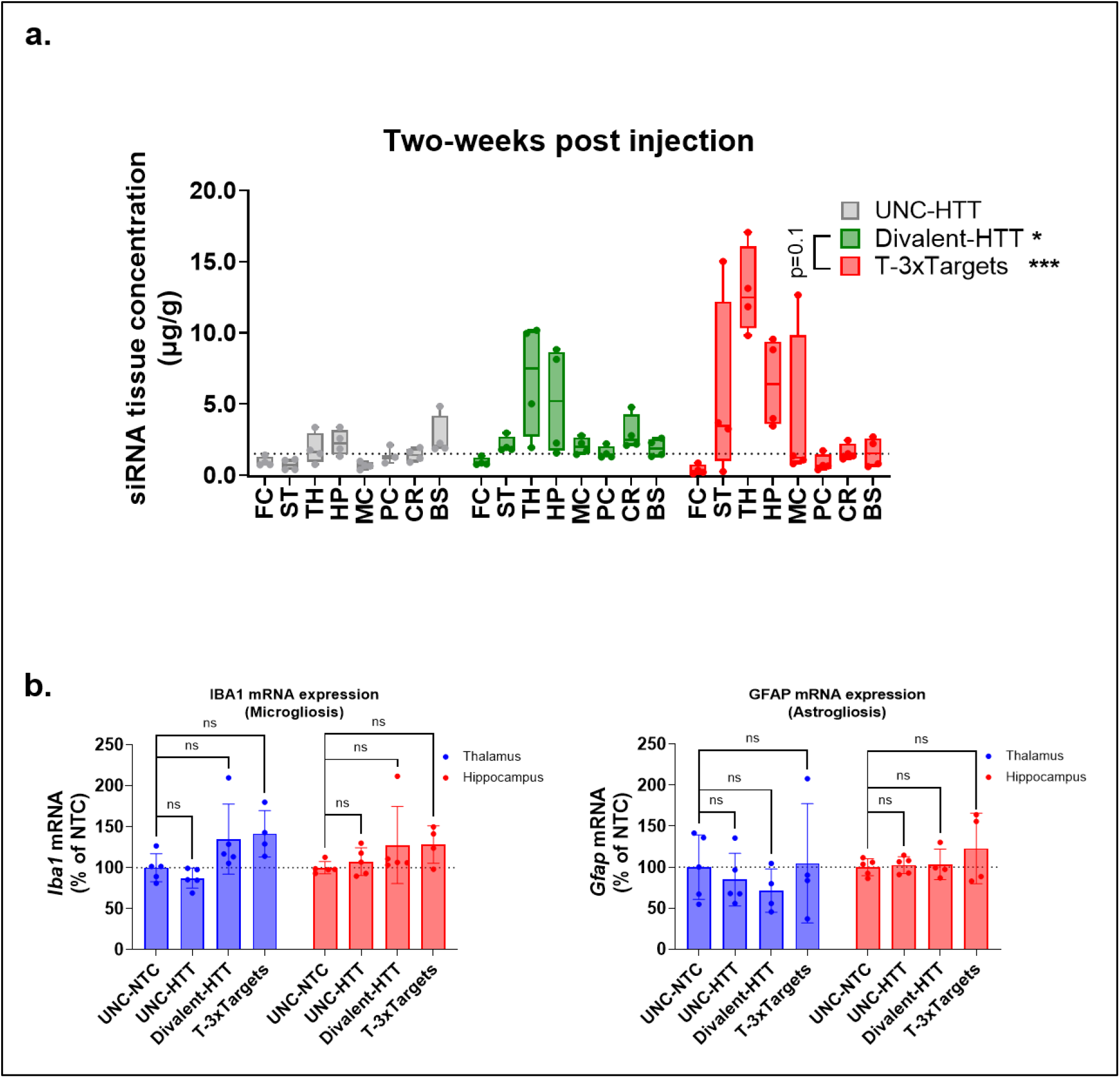
T-FMRN shows improved siRNA accumulation in the brain with a no apparent CNS toxicity. **a.** siRNA tissue accumulation two-weeks post intracereberoventricular (ICV) injection of UNC-HTT, Divalent HTT and T-3xTargets across various brain regions. **b.** mRNA levels of IBA1 and GFAP at two-months post ICV injection as markers of microgliosis and astrogliosisAll study groups are *n* = 4-5 mice/group. Statistical analysis was performed in GraphPad Prism: two-way ANOVA analysis followed by Tukey’s multiple comparisons across all groups for siRNA accumulation, and two-way ANOVA analysis followed by Dunnet’s multiple comparisons comparing to UNC-NTC control. (tissue accumulation measured by PNA assay; mRNA measured by the Quantigene 2.0 Assay; data represented as mean ± S.D.)

